# The diel disconnect between cell growth and division in *Aureococcus* is interrupted by giant virus infection

**DOI:** 10.1101/2024.05.01.592014

**Authors:** Alexander R. Truchon, Emily E. Chase, Ashton R. Stark, Steven W. Wilhelm

## Abstract

Viruses of eukaryotic algae have become an important topic due to their roles in nutrient cycling and top-down control of algal blooms. Omics-based studies have identified a boon of genomic and transcriptional potential among the *Nucleocytoviricota*, a phylum of large dsDNA viruses which have been shown to infect algal and non-algal eukaryotes. However, little is still understood regarding the infection cycle of these viruses, particularly in how they take over a metabolically active host and convert it into a virocell state. Of particular interest are the roles light and the diel cycle in virocell development. Yet despite such a large proportion of *Nucleocytoviricota* infecting phototrophs, little work has been done to tie infection dynamics to the presence, and absence, of light. Here, we examine the role of the diel cycle on the physiological and transcriptional state of the pelagophyte *Aureococcus anophagefferens* while undergoing infection by *Kratosvirus quantuckense*. Our observations demonstrate how infection by the virus interrupts the diel growth and division of this cell line, and that infection further complicates the system by enhancing export of cell biomass.

## INTRODUCTION

Diel patterns in phytoplankton are common. Specific factors known to cycle on a diel basis (*i.e.,* diel periodicity) among phytoplankton include population density, biomass, community species composition, intracellular metabolism, resources (*e.g.,* nutrients, organic constituents, DNA concentration), enzymatic activity, and primary production (Prezelin, 1992). Early field-based studies acknowledged these diel cycles, and began investigating the influence that time-of-collection for sampling might have on aquatic plankton research (Maxwell and Mikihiko, 1957; Yentsch and Ryther, 1957; Shimada, 1958; Doty, 1959). These studies included the domestication of oceanic phytoplankton to artificial light for *in vitro* investigations (Holmes and Haxo, 1958). By the 1960s, green algae, dinoflagellates, and diatoms had all been observed to have diel patterns *in vitro* (Harding et al., 1981). The dinoflagellate *Gonyaulax polyedra* divides on a diel cycle during a five hour interval near the end of a dark cycle / start of a light cycle (when under a 12/12 light:dark condition), and photosynthetic capacity follows the same cycling (Hastings et al., 1961). Generally, this cycle is thought to be decoupled from ambient light (Harding et al., 1981; Yoshikawa and Furuya, 2006), and an awareness of this cycle during experimental design enhances the reproducibility of results. For example, it was found that in phytoplankton collected from Sagami Bay (Japan), photosynthetic maximum (normalized absorption; mol C m^−2^ h^−1^) was highest at noon, and lower at dawn and dusk, with end-of-day timepoints being significantly variable (Yoshikawa and Furuya, 2006). Thus, photosynthetic variations were thought to be endogenously regulated (Behrenfeld et al., 2004). In the polymorphic haptophyte *Phaeocystis pouchetii,* it has been shown that synchronized cell division occurs midway through the dark cycle (on a 12/12 light:dark cycle), and that at higher light intensities or longer light cycles, cells could experience a division rate of greater than one per day (Jacobsen and Veldhuis, 2005). In a cell division periodicity study of 26 clonal cultures of marine algal cultures (representing 13 species), it was found that intraspecific variation does occur, and that different algal species can exhibit diel periodicity with division occurring at night or during the day (Nelson and Brand, 1979).

Recent examinations of diel periodicity have focused on the functional roles and dynamics of important marine microbes, which has been given further context through new methods including metatranscriptomics and flow cytometry (Aylward et al., 2015; Hu et al., 2018; Henderikx Freitas et al., 2020). One study measured metabolic activities of microbial life in the North Pacific Subtropical Gyre: metabolism was heightened during the day and related to carbon fixation and photosynthesis of autotrophs (Hu et al., 2018). The authors found that diel cycles in picocyanobacteria (*e.g., Synechococcus* and *Prochlorococcus*) included an increase in cell-size during the day, and cell division around dusk. Other studies have shown that picocyanobacterial gene expression is tied to diel periodicity (Zinser et al., 2009). However, picocyanobacterial cell counts remained stable in the gyre, as their diel metabolic activity was linked with the diel activity of dinoflagellates, haptophytes, ciliates and marine stramenopiles which would graze on the newly divided cells around dusk. Although viruses were not explored in that study, it is logical that they would also be influenced by their hosts reacting to the diel cycle.

In the North Pacific Subtropical Gyre (Aylward et al., 2015), diel periodicity of dominant photoautotrophs (*e.g., Ostreococcus* and *Prochlorococcus*) shapes community dynamics *via* light-based carbon acquisition at the base of the food web (Poretsky et al., 2009; Aylward et al., 2015). Collectively studies have explored the implications of diel periodicity in Bacteria, Eukaryota, and Archaea in important coastal and open ocean systems (Ottesen et al., 2014; Groussman et al., 2021; Muratore et al., 2022). However, there have been few studies exploring important bloom producing algae, nor how viral infection of algae and the formation of a virocell (a cell actively undergoing viral infection and thus with altered metabolic function) are affected by diel cycling.

In theory, the dominant algae within a system can change during blooms, establishing a new community dynamic still potentially coupled to diel cycles. Such effects could include shifts in populations grazing on a phytoplankton, shifts in primary productivity, and/or changes in light penetration of the water system. Viruses have been demonstrated to be important factor in bloom dynamics and specifically implicated in bloom termination (Jacquet et al., 2002; Brussaard et al., 2005; Steffen et al., 2017). Their role also has implications for biogeochemical cycling, including open ocean impacts on carbon cycling (*i.e.,* the viral shunt; Wilhelm and Suttle, 1999) and carbon export (*i.e.,* the viral shuttle; Sullivan et al., 2017). Indeed, a recent mesocosm study of carbon release showed viruses drove a 2-to 4-fold increase in extracellular carbon during bloom termination (Vincent et al., 2023). The effects of bloom events and bloom termination in the context of diel periodicity are important, especially given the diversity of life cycle strategies used by the causative agents of blooms and their viruses.

For the past two decades the brown tide bloom agent *Aureococcus anophagefferens* and “giant virus” *Kratosvirus quantuckense* (family *Schizomimiviridae*) have been studied in detail (Sieburth et al., 1988; Rowe et al., 2008; Truchon et al., 2023). The pelagophyte *A. anophagefferens* was characterized in 1985 (Sieburth et al., 1988) and has continued to produce blooms along the East Coast of the United States (Narragansett Bay, Barnegat Bay, Long Island bays) (Bricelj and Lonsdale, 1997), off the coast of China (near Qinhuangdao); Bohai Sea) (Zhang et al., 2012) and a bay on the South West coast of South Africa (Saldanha Bay) (Probyn et al., 2010). Brown tides are designated as harmful algal blooms (HABs) because of their economic and ecological detriment (Gobler et al., 2005). The virus *K. quantuckense* has been implicated as a regulator of *A. anophagefferens* brown tide bloom termination *via* population-wide cell mortality (Gastrich et al., 2004). Notably, irradiance levels have previously been tied to viral burst size in an *in vitro* setting, demonstrating virus particles produced during an infection cycle are dependent on the availability of light (Gann et al., 2020a). Given the need to better understand the physiological ecology and energetics of brown tides, we monitored the diel periodicity of this brown tide agent alone and during viral infection in lab studies. We observed a strict partitioning of physiological and metabolic processes by *A. anophagefferens* in relation to diel periodicity that was interrupted by viral infection.

## METHODS & MATERIALS

### Culture conditions

Three non-axenic isolates of *Aureococcus anophagefferens* were studied, including two that are resistant (strains CCMP1850 and CCMP1707) and a third (*A. anophagefferens* CCMP1984) susceptible to lytic infection by *Kratosvirus quantuckense*. Cultures were maintained at 19° C under a 12:12 light dark cycle in ASP12A growth media (Gann, 2016). Light levels for maintenance and experimental cultures were ∼70 µmol photons m^-2^ s^-1^. Shading experiments were conducted by wrapping one or two layers of neutral density screening around individual culture tubes that reduced irradiance to 40 and 20 µmol photons m^-2^ s^-1^, respectfully. Prior to experimentation under reduced light conditions, cultures were moved and acclimated to the specific light treatment for at least 72 h. The concentration of cells in *A. anophagefferens* cultures was determined using a CytoFLEX flow cytometer (Beckman Coulter, Brea, CA) (Chase et al., 2022). Abundance in samples for gated cellular populations was quantified *via* violet side scatter (V-SSc) versus chlorophyll fluorescence (absorption 488 nm, emission 690 nm) (Chase et al., 2022).

### Cell diameter measurements

Individual cell diameters were determined based on measurements from a FlowCam 8000 (Fluid Imaging Technologies, Scarborough, ME). Briefly, culture samples were diluted to approximately 1 x 10^6^ cells/mL and then imaged using the FlowCam’s 20X objective. Ten thousand individual cellular images were used to calculate average cell diameter and volume as well as for verification of cell concentration (using VisualSpreadsheet 2). FlowCam measurements for average diameter were compared to flow cytometry estimates (CytoFLEX C07821). *A. anophagefferens* CCMP1984 was compared on both devices over the course of 24 h after either being infected (see below) or not infected with *Kratosvirus quantuckense*. To determine the relation between these measurements, a Pearson’s coefficient was calculated.

### Infection with Kratosvirus quantuckense

*A. K. quantuckense* particles were generated by infection of 1 L of a 7d-old culture of *A. anophagefferens* CCMP1984 grown in ASP_12_A medium as above. After allowing the population to lyse (14 d), lysate was filtered sequentially through 1-µm and 0.45-µm pore-size, 47-mm diameter low protein binding Durapore (PVDF) membrane filters (MilliporeSigma; Burlington, MA). Viruses in the filtered lysate were concentrated *via* tangential flow filtration through a 30 kDa Pelicon XL (MilliporeSigma; Burlington MA) filter to an approximate volume of 50 mL as previously described (Coy and Wilhelm, 2020). Following concentration of viruses from lysate, contaminating bacteria were removed *via* centrifugation (3,500 xG, 10 min). Viral particles were enumerated *via* flow cytometry (Chase et al., 2023). Briefly, lysate was fixed with 1% glutaraldehyde solution in the dark at 4° C for at least 1 h. Lysate was then stained with SYBR Gold (Invitrogen; Waltham, MA) at a final concentration 0.5X at 80° C for 10 min. Virus particles were enumerated on a CytoFLEX flow cytometer (C07821) equipped with a violet and a blue laser (Beckman Coulter; Brea, CA) (Chase et al., 2023; Zhao et al., 2023).

For experiments, infection of *A. anophagefferens* was performed on cells in exponential growth stage diluted to 1 x 10^6^ cells ml^-1^ in fresh ASP_12_A medium. Viral lysate was added to diluted *A. anophagefferens* cells at a multiplicity of infection (MOI) of ∼100 viral particles per *A. anophagefferens* cell (unless otherwise specified) to approach uniform infection (Gann et al., 2020a). Cell concentrations during infection were determined *via* flow cytometry (Chase et al., 2022). Following lysis of samples, aliquots were fixed using 1% glutaraldehyde for further enumeration of released viral particles. When infecting at lower MOIs, cell diameter measurements were determined *via* flow cytometry 23 hr following infection before the initiation of the light cycle. This allowed for measurements of cell size before cell lysis without the input of any additional light.

### Transcriptome analyses of A. anophagefferens infection by Kratosvirus quantuckense

We took advantage of an existing transcriptomics data set (Moniruzzaman et al., 2018) to query the progression of infection at the molecular level. Trimmed reads of infected and uninfected cultures of *A. anophagefferens* were analyzed using DESeq2 in R (Love et al., 2014). Control (uninfected) samples were compared to identify shifts in gene expression throughout the diel cycle. Control time points were divided into four periods based on when samples were collected during the original transcriptome. These samples were defined as early day, taken between two and three hours after the initiation of the light cycle, late day, taken eight hours after the initiation of the light cycle, early night, taken approximately 30 min after the initiation of the dark cycle, and late night, taken nine hours after the initiation of the dark cycle. Individual genes with a log_2_-fold change of at least 2 and a p-value of < 0.05 for at least two of the four periods were defined as differentially expressed. The same parameters were applied to identify differentially expressed genes between control and infected treatments, though due to a limited number of identifiable genes at this level a log_2_-fold change of > 1.5 was used to identify other potentially altered expression levels. To identify other genes of interest that may be up or downregulated at a specific timepoint at a lower significance level, a log_2_-fold change of 0.58 (a fold change of > 1.5) was also examined. As a caveat, downregulation and upregulation will be used to equate proportional representation of mapped reads between treatments throughout this paper.

To examine infection-driven inhibition of cell division, differentially expressed genes were filtered to only those associated with the cell cycle based on functional annotation in the *Kyoto Encyclopedia of Genes and Genomes* (KEGG) pathways map04110 (Cell cycle), map04111(Cell cycle – yeast), map04210 (Apoptosis), map04115 (P53 signaling pathway), and map04218 (Cellular senescence). Read abundance calculations for individual gene transcription trends and for incorporations into heatmaps were performed using the transcripts-per-million (TPM) method (Wagner et al., 2012). Heatmaps were constructed using *Heatmapper.ca* (Babicki et al., 2016) with genes clustered *via* single linkage clustering.

### Determination of vertical transport rates

To assess the sinking rate of *A. anophagefferens* CCMP1984, cells in logarithmic growth were inoculated into a vertical settling column (**Figure S1**) containing 225 mL ASP_12_A and allowed to settle. After 2 h the bottom 50 mL of the column was drained through the collection tube and agitated to homogenize the cells to a uniform concentration. This process was repeated for the remaining 200 mL of media. Cell concentrations and diameters were calculated *via* flow cytometry. To determine the sinking rate of infected cells, *A. anophagefferens* CCMP1984 was infected with *K. quantuckense* at an MOI of 100 either 2 h (early infection) or 16 h prior (late infection) to sinking rate assessment. Sinking velocity (Ψ) was calculated using the following formula:

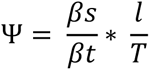

In which *βs* is total exported cells, *βt* is total cells in the column, *l* is distance traveled in meters, and *T* is time in hours (Bienfang, 1981; Mao et al., 2021).

### Statistical Analyses

Statistics were performed using Prism 9.1.0. Differences between growth rates, cell sizes, and sinking rates were determined using a one-way ANOVA followed by Tukey’s *post hoc* testing. Correlation coefficients were determined using a simple linear regression. Non-metric multidimensional scaling (nMDS) and hierarchical clustering analysis of host transcriptional shifts in the uninfected reference transcriptome was performed in PRIMER v7.0 (Clarke, 2015) using a Bray-Curtis dissimilarity matrix.

## RESULTS

### Diel partitioning of metabolic functions

*A. anophagefferens* CCMP1984 cultures grown under a 12:12 light:dark cycle were observed every 4 h during the light period to determine cell concentration and relative fluorescence measurements throughout the light cycle. *A. anophagefferens* cell diameter determined from the FlowCam 8000 was correlated strongly with the violet side-scatter (V-SSc) measurements detected using a 405 nm violet laser (**p < 0.001; R^2^ = 0.9413**) (**Figure S2**). This correlation was consistent for both virus-infected and uninfected *A. anophagefferens* cells through different stages of the growth and infection cycle. For this reason, V-SSc was used as a proxy for average cell diameter for the remainder of the experimentation.

During light periods, *A. anophagefferens* CCMP1984 cell densities were generally constant (**Figure 1A**). However, after the dark period cell abundance increased, consistent with a diel association with cellular division. V-SSc relative fluorescence indicated a similar pattern between the light and the dark cycle, with average cell diameter (**Figure S2**) and V-SSc fluorescence (**Figure 1B**) increasing throughout the day and reducing during the night.

**Figure 1.**
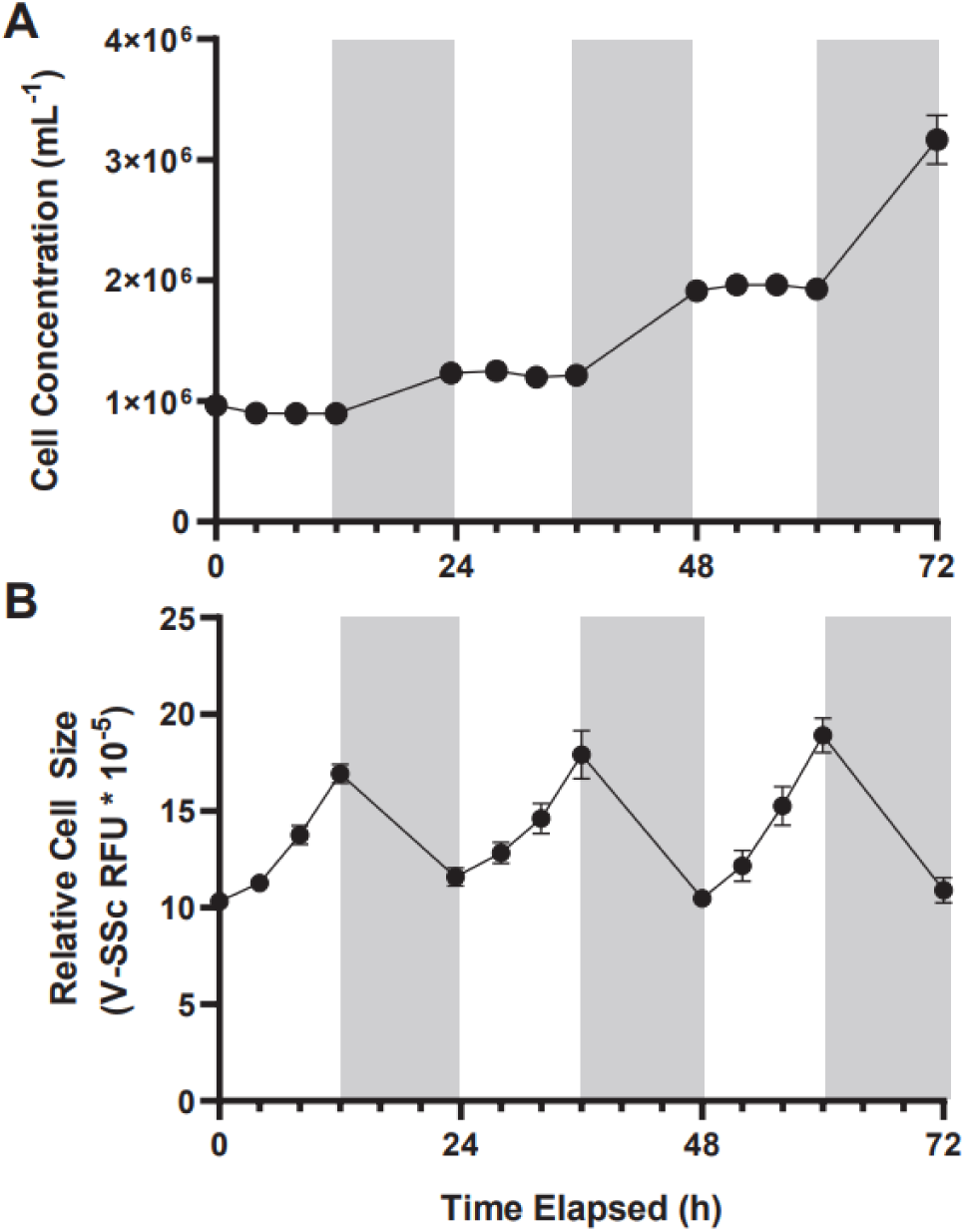
Cell concentration (A) and average cell diameter as measured *via* violet side-scatter (B) of *A. anophagefferens* over the course of 72 hours with samples taken every four hours during the light cycle. Periods of light are indicated in white, and periods of dark are indicated in gray (n = 5).

Population-wide increases in cell size were not linear, with the rate of cell diameter growth increasing along with the length of exposure to light. While other strains of *A. anophagefferens* differed in specific growth rate and percent change in cell size over light and dark periods, all strains we tested followed the pattern of division in the dark, impeded division during the day, and cyclical cell-size changes (**Table 1**). A significant increase in cell density occurred within eight hours of the dark period for CCMP1984 (p = 0.024) and within 12 h of the dark period for CCMP1850 (p = 0.051) (**Figure S3**).

**Table 1.** Mean growth rate and changes in cell diameter through either light or dark periods of three different strains of *A. anophagefferens*. Standard deviation is denoted in parentheses.

|  | CCMP1850 | CCMP1984 | CCMP1707 |
| --- | --- | --- | --- |
| Total Growth Rate ($D^{-1}$) | 0.373 ($\pm 0.06$) | 0.386 ($\pm 0.05$) | 0.251 ($\pm 0.04$) |
| Day Growth Rate ($D^{-1}$) | 0.089 ($\pm 0.07$) | 0.023 ($\pm 0.05$) | 0.026 ($\pm 0.05$) |
| Night Growth Rate ($D^{-1}$) | 0.657 ($\pm 0.13$) | 0.749 ($\pm 0.12$) | 0.475 ($\pm 0.06$) |
| Day Change in Cell Diameter (%) | 23.44 ($\pm 2.58$) | 35.54 ($\pm 2.79$) | 15.15 ($\pm 1.36$) |
| Night Change in Cell Diameter (%) | -26.25 ($\pm 3.03$) | -30.35 ($\pm 3.68$) | -18.96 ($\pm 1.25$) |

Given *A. anophagefferens* CCMP1984 has been maintained in our laboratory for an extended period (over 10 years) and thus may have grown accustomed to these light settings, circadian rhythms could not be ruled out as a factor in diel-associated cellular growth and cell division. To test this possibility, *A. anophagefferens* cultures were exposed to reduced light levels. While no discernable growth differences were detected between different light levels (p-value = 0.213), cell diameters in cultures maintained at 20 µmols photons m^-2^ s^-1^ had significantly (p-value < 0.0001) smaller diameters (∼ 2.4 µm), as compared to high light treatments (2.9 µm). Cells in low light cultures increased in diameter by 39.3%, while full irradiance cultures increased in diameter by 114.8% (**Figure 2A-C**). Population growth was also significantly impeded in low light cultures over 72 h (**Figure 2D**). Entraining cultures on a 12:12 light dark cycle only to either leave the lights on after 12 h of light or leave the lights off after 12 h of darkness showed that *A. anophagefferens* did not in these instances display any characteristics of a free-running clock (**Figure S4**). Furthermore, *A. anophagefferens* had growth rates of approximately zero or lower when exposed to 24-h light or 24-h dark (**Table S1**). In 24-h light, cells continuously increased in size over the course of 48 h, while cells continuously decreased in size in 24-h darkness (**Table S1**). While increasing the length of the light period did lead to continued increases in cell size, longer light periods did not have much effect on net growth rate.

**Figure 2.**
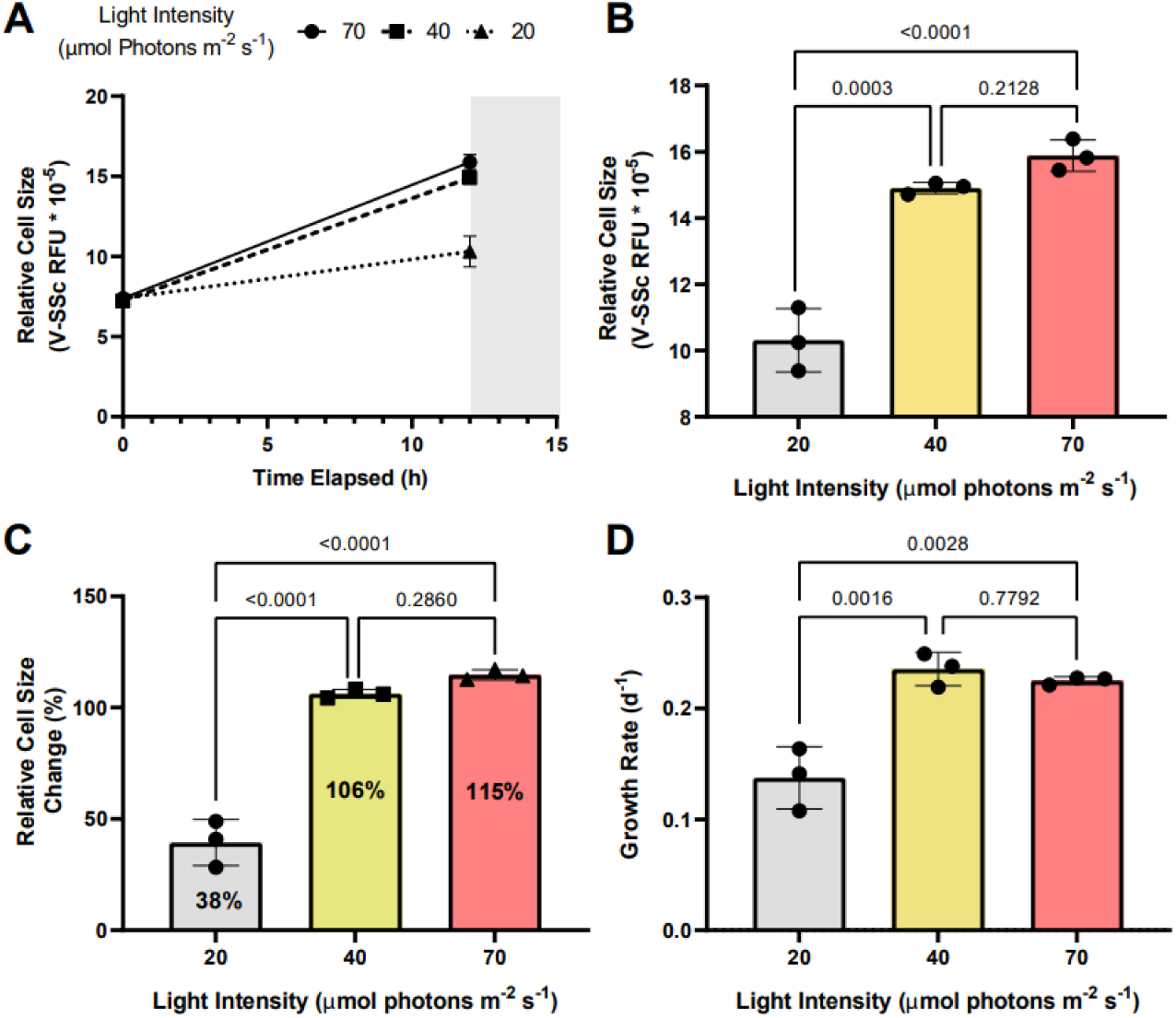
Change in average *A. anophagefferens* CCMP1984 cell diameter as measured *via* violet side-scatter over the course of a single 12-hour light period (A) and final average cell diameter at the termination of the light period (B) with cultures kept at 3 different irradiance levels (high: red, ∼70 µmol m^-2^ s^-1^; medium: yellow, ∼40 µmol m^-2^ s^-1^; low: grey, ∼20 µmol m^-2^ s^-1^). (C) Growth rate of *A. anophagefferens* over the course of 72 h under the 3 irradiance levels. P values are represented above separate irradiance levels compared *via* two-way ANOVA and post-hoc multiple comparisons adjusted with Tukey’s HSD (n = 3).

### Infected cells increase in diameter while division is inhibited

*A. anophagefferens* CCMP1984 was infected with *K. quantuckense* to observe the effects of viral infection on the diel growth cycle. During the first 12 h of infection, (during the light cycle) no differences were observed between infected and control samples treated with filtered lysate (**Figure 3**). However, during the night cycle, infected cultures did not divide and stayed at the same cell concentration (**Figure 3A**) and cell diameter observed at the termination of the light cycle (**Figure 3B**). Following the first 24 h, average cell diameter increased again, up to additional 21% increase from the first light cycle (**Figure 4C**). It is unclear if the cells that continued to increase in size were the same group of infected cells or were instead previously uninfected cells that are continuing to skew the average size higher as they continue to grow. To determine whether size shifts between infected and uninfected cells were evident in a single culture, size was measured at lower MOIs approximately 23 h following infection. A higher MOI led to an increased average diameter of infected cultures following the night cycle, with cultures infected at an MOI of 100 displaying a 30% increase in average V-SSc as compared to those infected at an MOI of 10 (**Table 2, Figure S5**). A negative correlation (R^2^ = 0.9427, slope = - 0.372) between MOI and percent similarity in size of infected cultures to control cultures was found (**Figure S6A**).

**Figure 3.**
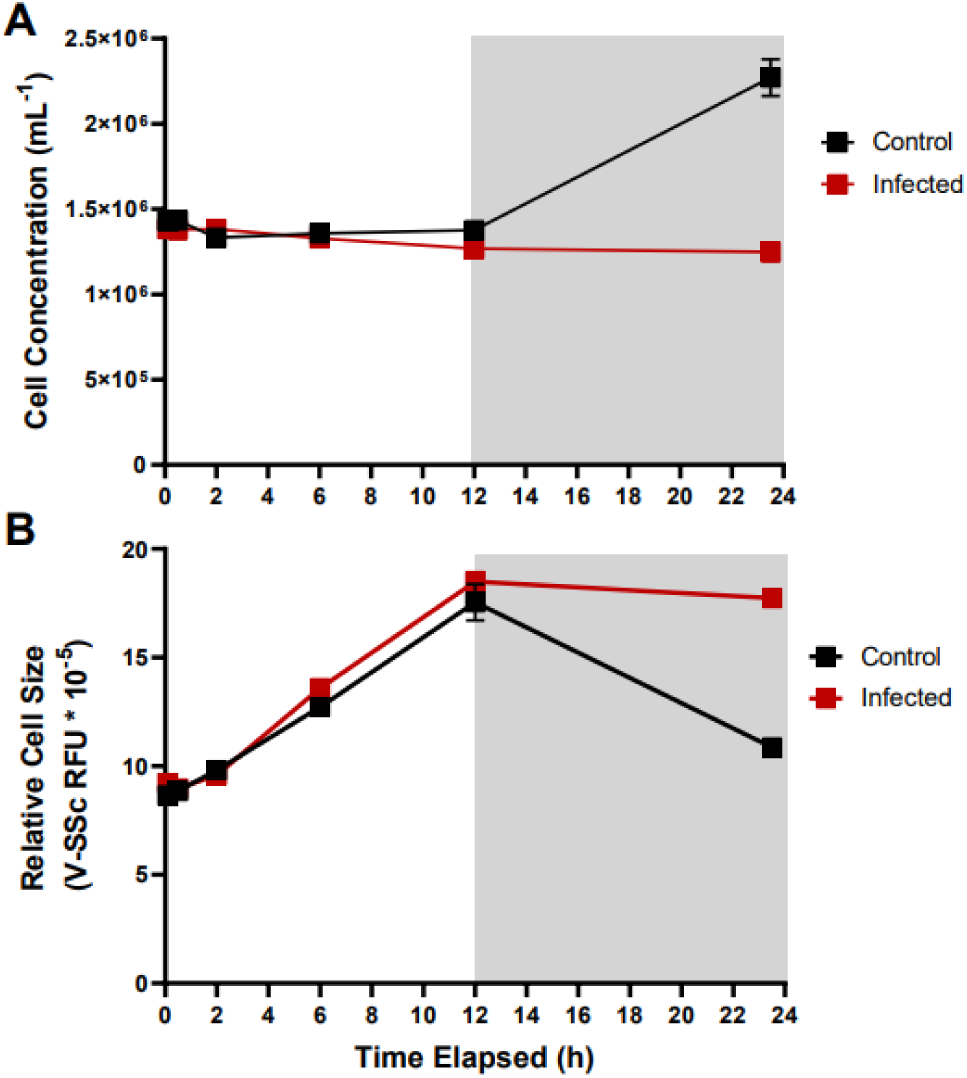
Cell concentration (A) and average cell diameter as measured *via* violet side-scatter (B) of *A. anophagefferens* over the course of 24 hours in the presence of *K. quantuckense*. Samples treated with viral lysate are indicated in red and control samples are indicated in black. Light periods are indicated by a white background while dark periods are indicated with a grey background (n = 3).

**Figure 4.**
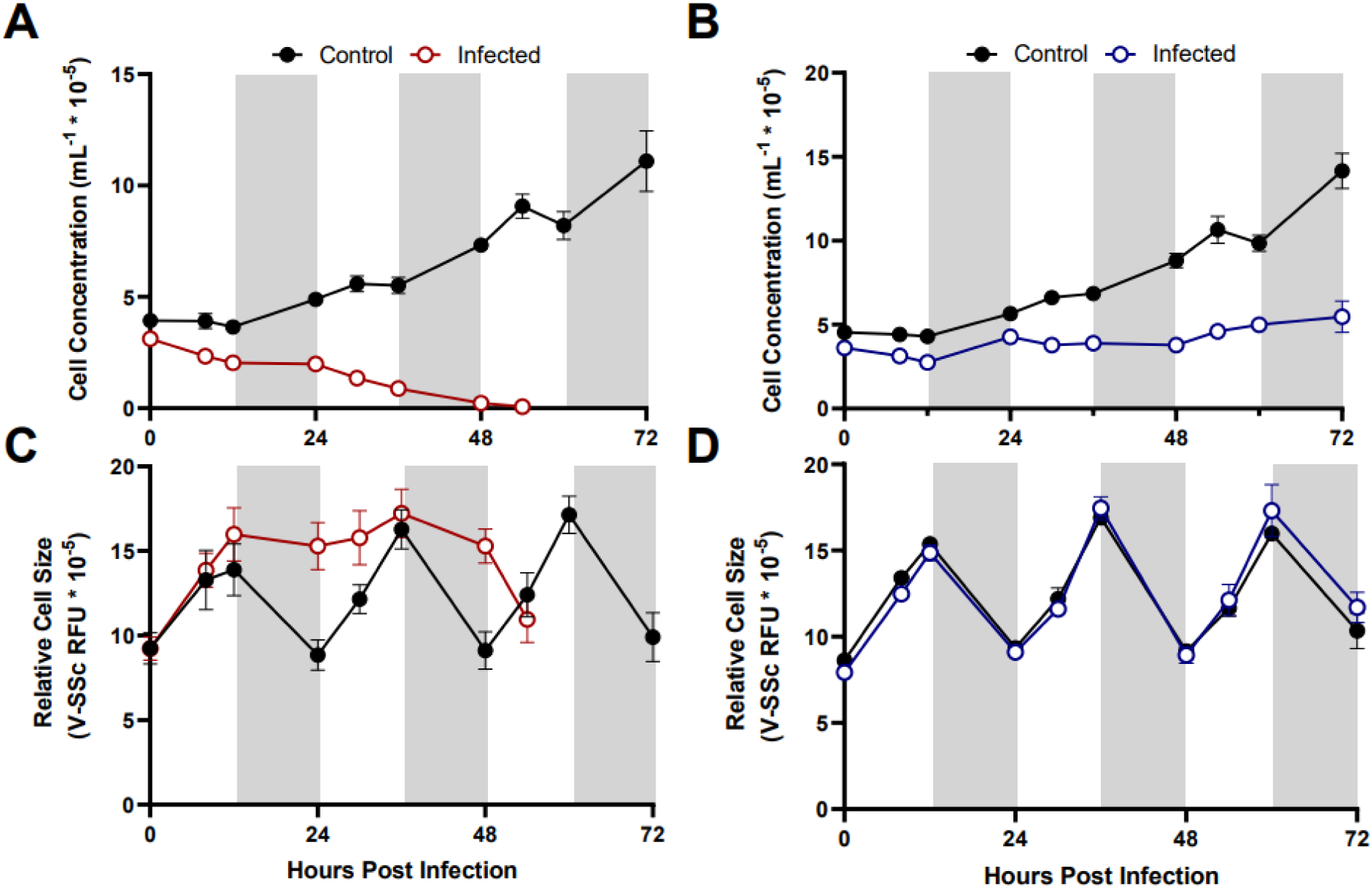
Growth of different *A. anophagefferens* strains CCMP1984 (A, C) and CCMP1850 (B, D) in the presence and absence of viral lysate over the course of 72 hours. Cell concentrations (A-B) and relative cell size (C-D) are denoted. Light periods are indicated by white backgrounds while dark periods are indicated by gray backgrounds. Black lines represent uninfected cultures, solid colored lines represent cultures infected at the initiation of the light cycle (n = 3).

**Table 2.**
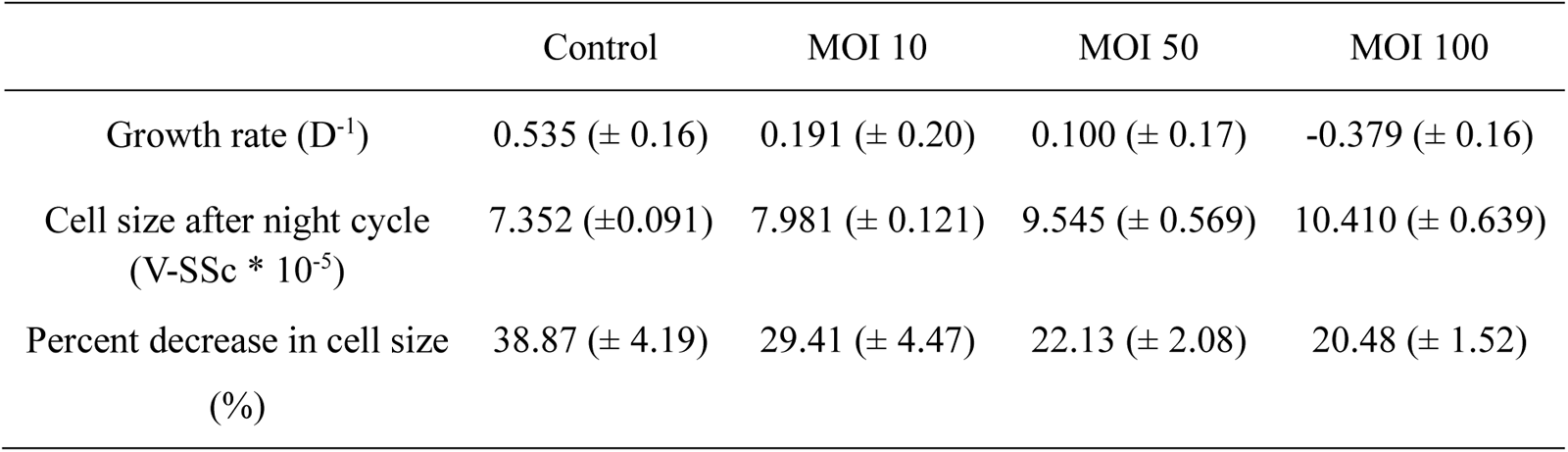
Growth and division characteristics of *A. anophagefferens* CCMP1984 23 hours following viral infection with *K. quantuckense* at variable MOIs. Cell size following the night cycle was recorded prior to the initiation of light cycle. Percent change in cell size was determined based on the peak cell size at the initiation of the dark cycle and the final size at the end of the dark cycle. Standard deviation is denoted in parentheses.

Another *A. anophagefferens* strain which displayed a resistant phenotype to viral infection by *K. quantuckense* was tested for their physiological response to viral exposure over the course of several days. Continued growth following the initial 24-h period in infected strains was evident in CCMP1984, before average cell diameter diminished coinciding with the total lysis of the culture around 48 hours (**Figure 4**). CCMP1850 also appeared to be inhibited in cell division (**Figure 4B**) but maintained normal cell size cycling throughout the light:dark period as compared to uninfected cells (**Figure 4D**). This may imply that whatever stress viral presence places on this strain, it is unrelated to diel cycling and the cell cycle.

### Separate cell cycle transcriptomic activity between day and night

We returned to a transcriptome from a previous infection study (Moniruzzaman et al., 2018) to identify cell division genes differentially expressed between the light and dark periods. Samples were subdivided into four categories: the early day (n = 9, two to three hrs. into the light period), late day (n = 3, eight hrs. into the light period), early night (n = 3, ∼30 min. into the dark period), and late night (n = 3, nine hrs. into the dark period). Cluster analyses revealed an almost cyclical relationship among uninfected samples with similarity between categories strongest for neighboring groups (*e.g.,* late day was most like early day and early night) (**Figure S7**). Although we will focus on cell cycle-associated genes, in total 1,823 genes were identified as differentially expressed between at least two of the time periods analyzed under our highly conservative parameters.

A clear partition in cell cycle gene expression throughout different stages of the day was evident (**Figure 5**). Periods that differed most notably from one another were the early day/early night (22 differentially expressed genes; **Figure S8**) and the late day/late night (13 differentially expressed genes; **Figure S9**). Cohesin subunit genes (*scc1, scc2, scc3, smc1,* and *smc3*) were differentially expressed between early morning and early night (**Figure 5; Figure S8**) with a consistent drop off in expression during the late night and markedly low expression throughout the day (**Figure 5; Figure S9**). Condensin subunits *smc4*, and *ycs4* were overexpressed during the late-night timepoint as compared to the day, with steady down regulation of *ycs4* in the early night (**Figure 5, Figure S8, Figure S9**).

**Figure 5.**
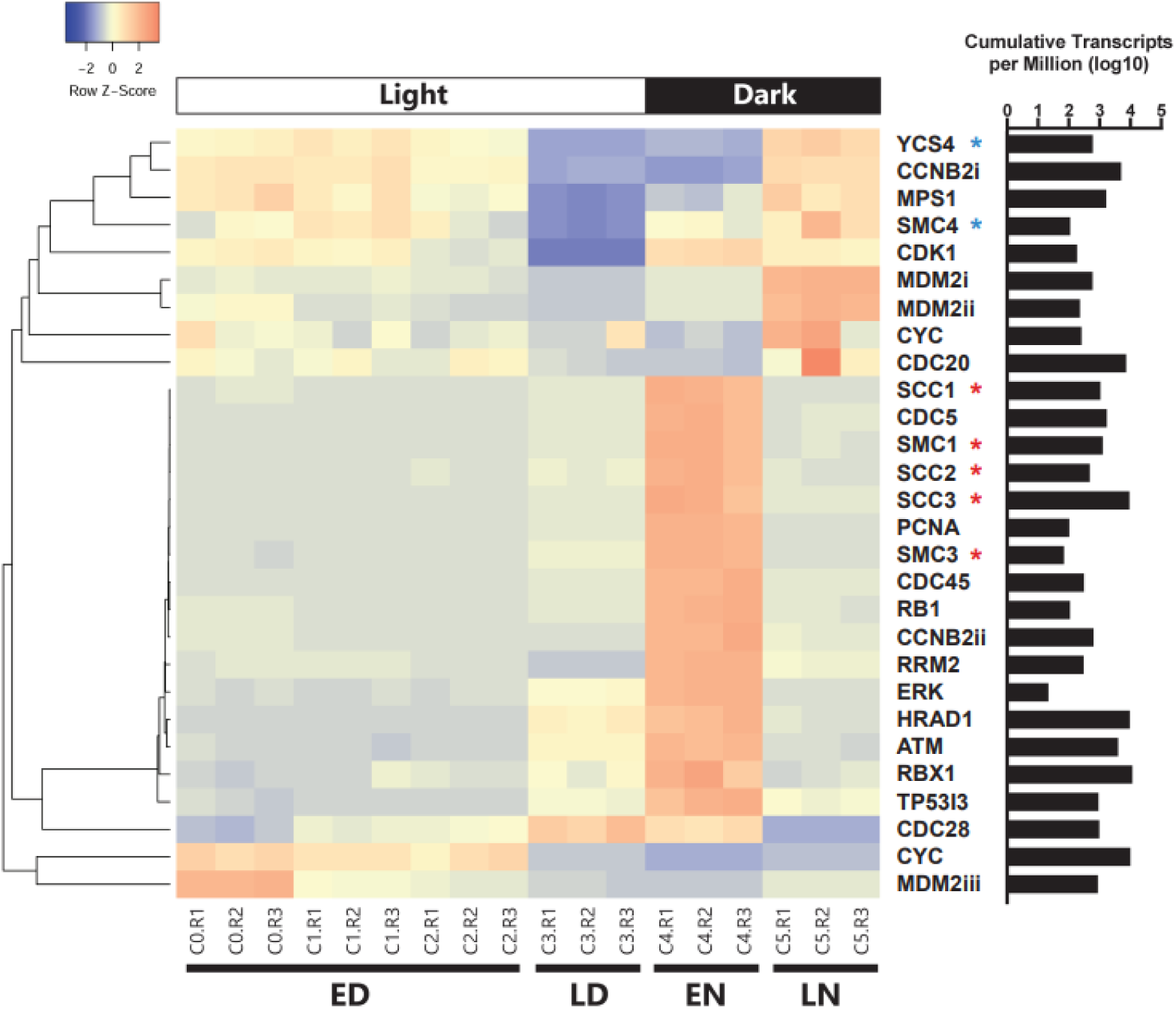
Clustered heatmap containing all cell cycle genes based on KEGG databases that were differentially expressed between at least two of the four categorical time points (ED: early day; LD: late day; EN: early night; LN: late night). Rows were clustered through Pearson correlation as indicated by the cladogram and z-scored based on TPM values. The sum of transcripts across all treatments for each individual gene are indicated in log(TPM).

Regarding cyclin associated genes and their expression throughout the cell cycle, *ccnb2* (Cyclin B) is expressed significantly more at night as compared to the day (log_2_ fold change = 2.47, p < 0.001), though one homolog (*ccnb2i*; 12473) is under expressed in the early night as compared to the morning. A *cdc20* and a *cdk1* homolog were expressed late at night, but not earlier. C*dc45* was also expressed early in the night and not present late at night.

A homolog for tumor (*i.e.,* cell division) suppressor gene p53 is not encoded by *A. anophagefferens*. Still, certain homologs of genes associated with p53 do display changes in expression. In the early day *mdm2* is highly expressed, while it is downregulated in the early night. It is also highly expressed late at night and barely expressed mid-day. A homolog of tumor suppressor *rb1* is differentially expressed in the late day/early night period from the morning. Cell cycle regulation genes *tp53i3* (tumor protein p53 inducible protein 3), *hrad1* (Rad1 checkpoint protein homolog), *atm* (serine/threonine kinase), *erk* (extracellular signal-regulated kinase), and *rrm2* (ribonucleoside reductase regulatory subunit M2), many of which are expressed downstream of p53, also follows this pattern (**Figure 5**).

### DNA damage and p53-associated genes respond to *Kratosvirus quantuckense* infection

Given cell division appears inhibited when *A. anophagefferens* is infected with *K. quantuckense*, we sought to identify associated genes that might be targeted by the virus for regulation. When comparing the infection transcriptome to the control, only 11 cell cycle associated genes were identified at a log_2_-fold change of ≥ 1.5 (**Table 3**). While most of these genes were in the late night time point of the infection cycle, three genes were identified 12 h following infection (early night) and one at the 6 h mark (late day). Interestingly, all three genes identified at the 12 h timepoint were homologs of the *mdm2* gene, two of which were down-regulated in the infected samples with the other highly up-regulated. While all these genes increase in expression as the night continues, their role in infected samples in the early night may also be relevant.

**Table 3.** *A. anophagefferens* cell cycle-associated genes that are differentially expressed (p-value < 0.05, log_2_fold change > 1.5 or < -1.5) at either the 6-h, 12-h, or 21-h timepoints between infected and uninfected samples.

| Gene ID # | Gene Name | Time | Direction | Fold Change<br>(log<sub>2</sub>) | Notes |
| --- | --- | --- | --- | --- | --- |
| 32157 | *cdc15* | 6 | Down | -1.51 | Protein kinase; cell division control protein |
| 12504 | *mdm2* | 12 | Up | 2.79 | E3 ubiquitin-protein ligase; p53 regulation |
| 24297 | *mdm2* | 12 | Down | -1.72 | E3 ubiquitin-protein ligase; p53 regulation |
| 3154 | *mdm2* | 12 | Down | -1.51 | E3 ubiquitin-protein ligase; p53 regulation |
| 36910 | *smc2* | 21 | Up | 1.84 | Structural maintenance of chromosome;<br>Condensin subunit |
| 58667 | *skp1* | 21 | Up | 1.82 | S-phase kinase-associated protein 1; Myt1<br>regulator |
| 70503 | *smc1* | 21 | Up | 1.81 | Structural maintenance of chromosome;<br>Cohesin subunit |
| 70163 | *pcna* | 21 | Up | 1.77 | Proliferating cell nuclear antigen |
| 72635 | *smc3* | 21 | Up | 1.55 | Structural maintenance of chromosome;<br>Cohesin subunit |
| 72516 | *myt1* | 21 | Down | -1.75 | Mitosis inhibitor protein kinase |
| 72033 | *smc4* | 21 | Down | -1.5 | Structural maintenance of chromosome;<br>Condensin subunit |

At the 21-h timepoint, seven cell cycle genes were differentially expressed between control and infected samples, including four SMC-like genes. Two cohesin-associated genes were up-regulated (*smc1* and *smc3*), one condensin-associated gene was up-regulated with the other down-regulated (*smc2* and *smc4*, respectively) (**Figure S10**). The cell cycle regulatory genes *skp1* and *pcna* were also upregulated at this time point (Liu et al., 2005; Zhang et al., 2023). Another regulator of the cell cycle, a homolog for the *myt1*/*wee1* gene which induces cell cycle delay, was downregulated (Détain et al., 2021).

Using a less conservative method for defining differential expression (*i.e.,* the 1.5-fold change described in Moniruzzaman *et al.,* 2018) an additional 64 cell cycle genes were differentially expressed between all infected and control samples (**Figure 6, Table S2**). Among these genes, seven were consistently upregulated in infected samples, 19 were consistently downregulated, and five alternated between upregulated and downregulated. Multiple inhibitors of cell cycle regulation genes (*myt1/*wee1) are consistently upregulated, including a protein arginine methyltransferase (*prmt5*) and *rbx1* which can target CDKIs (Jia et al., 2011; Beketova et al., 2022). Also upregulated was a ubiquitin-protein lyase homolog (*siah1*), which can act downstream from p53 in cell cycle arrest pathways (Frew et al., 2002).

**Figure 6.**
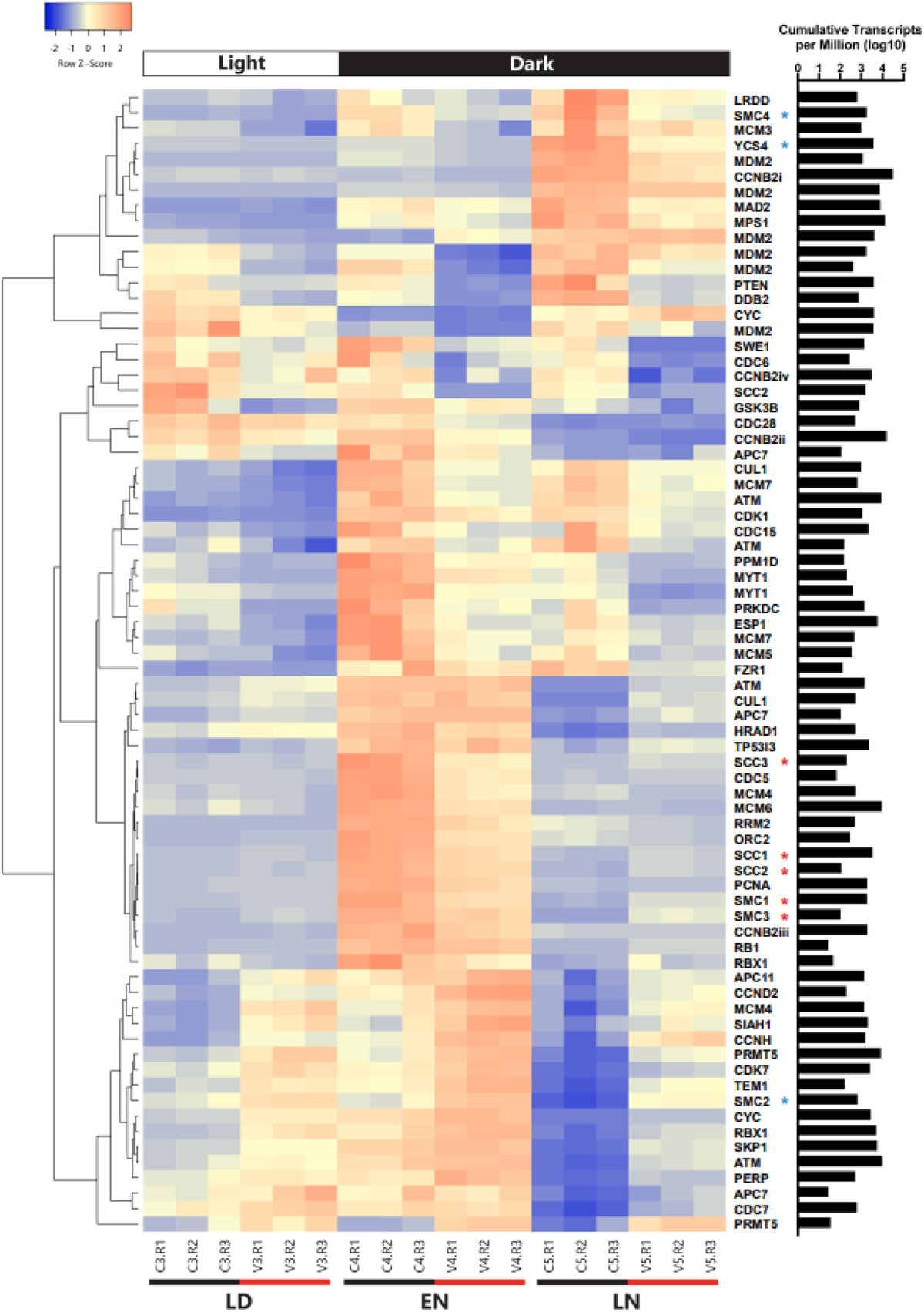
Clustered heatmap containing all cell cycle genes that were differentially expressed between infected and uninfected *A. anophagefferens* at one or more timepoints (LD: late day; EN: early night; LN: late night). Early morning timepoints were not included due to high variability in gene expression among infected samples at this period. Infected samples are indicated by a V and control samples are indicated by a C. Rows were clustered through Pearson correlation as indicated by the cladogram and z-scored based on TPM values. The sum of transcripts across all treatments for each individual gene are indicated in log(TPM).

Of the consistently down-regulated genes, many are associated with DNA replication and checkpoints for DNA damage repair. Regarding initiation of replication, two mini-chromosome maintenance (MCM) subunits and one origin recognition complex (ORC) subunit were downregulated in infected samples. Likewise, downregulated genes associated with suppressing the cell cycle due to DNA damage include *rrm2*, *prkdc*, *ddb2*, and *myt1/wee1* (**Figure S10**) (Chen et al., 2021a; Chen et al., 2021b; Détain et al., 2021; Zuo et al., 2024). Condensin and separase (*esp1*) genes decreased in expression as well. Outside of genes associated with DNA replication and repair are various other downstream effectors of the conventional p53 pathway. Negative feedback regulators including the previously mentioned three *mdm2* homologs and the *ppm1d* protein phosphatase were downregulated throughout the infection cycle.

In addition to shifts in expression across all time points, genes differentially expressed at single time points may provide insight into how viral infection impacts the cell cycle of *A. anophagefferens.* For instance, while several condensin-associated genes are downregulated at the 12-hour time point, they become upregulated at the 21-hour time point. Such dynamics implicate *K. quantuckense* derived regulation of host genes beyond simply turning a gene on or off. The *rb1* homolog was also upregulated at the 21-hour time point (Mendoza et al., 2014). Interestingly, while a cyclin B homolog was downregulated at the 21-hour time point, cyclin D and cyclin H were upregulated at 12 and 21 hpi, respectively (**Table S2**). Cyclin-dependent kinases also acted contrarily, with some (*cdc28*, *cdc6*, *cdc5*, *cdc15* and *cdk1*) downregulated at certain time points and other (*cdc7* and *cdk7*) upregulated (**Table S2**).

### Infection alters host cell sinking rate

To assess biophysical consequences of diel shifts in cell size / composition in the presence and absence of *K. quantuckense*, sinking rates of *A. anophagefferens* were measured. In calm waters (no upwelling) *A. anophagefferens* sinks in the water column and does not maintain a natural buoyancy. The sinking velocity of uninfected *A. anophagefferens* cells was constant regardless of time of day or cell size (**Figure S11**). Within two hours of infection of *A. anophagefferens by K. quantuckense*, the alga sinking rate increased compared to control (uninfected) cells (**Figure 7A**). A similar trend was noted 16 h after infection, suggesting cells in the early and late stages of the virocell state would be exported from the water column at the same rate (**Figure 7A**). When comparing cell diameter or volume for uninfected cells at the bottom of the settling column to cells at the top, there were no significant differences (**Figure 7B-C**).

**Figure 7.**
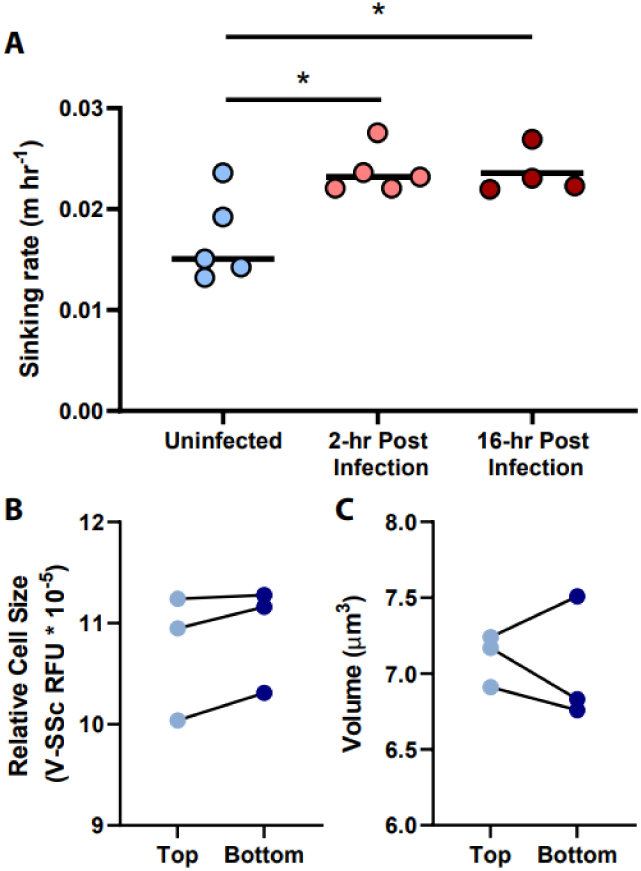
Sinking rate of *A. anophagefferens* CCMP1984 cells as it relates to the length of infection (A). Relative fluorescence as a proxy for cell diameter (B) and volume as determined *via* the FlowCam (C) of CCMP1984 in the top portion and bottom portion of the sinking column after incubation.

## Discussion

In the past, studies of marine algal growth patterns have generally been conducted with daily sampling at consistent time points (relative to light:dark cycles) (Tang, 2003; Shirai et al., 2008; Perrin et al., 2016; Gann et al., 2020b; Gann et al., 2022). While this method increases comparability over long-term sampling schemes, it excludes physiological changes that occur in response to prolonged exposure to light or the absence of light. Thus, other time points (*i.e.,* 6 daylight hours, 12 daylight hours, *etc.*) likely need to be considered as they potentially offer other physiological states. Furthermore, studies that have focused on hourly changes in algal growth dynamics often have not considered the growth cycle, physiology, and metabolism of virocells which may make up a significant portion of the natural community (Brussaard et al., 1996; Vincent et al., 2021). Indeed at least some algal viruses rely on light during infection (Derelle et al., 2018; Gann et al., 2020a) with certain giant viruses even encoding rhodopsins (Needham et al., 2019). Collectively this implicates the diel cycle as a potential modulator of virus activity in phototrophs. We examined physiological shifts of infected and uninfected *A. anophagefferens* in the context of diel periodicity. We also explored transcriptomic data to understand shifts in gene expression in infected and uninfected cells.

### Cellular and population growth of *A. anophagefferens* is constrained by the diel cycle

*A. anophagefferens* cell diameter gradually increased during the light period, with cell division (*i.e.,* size reduction and cell density increase) almost exclusively occurring during the dark period. While this separation of growth and division has been observed in phytoplankton, rarely has the distinction been so distinct, as cellular growth is often seen during both dark and light periods (Harding et al., 1981; Goto and Johnson, 1995; Jacobsen and Veldhuis, 2005; Moulager et al., 2007). Moreover, we observed no change in cell concentrations during the light period, with occasional (yet statistically insignificant) decreases in cell count when the light period is increased (**Table S1**). Attempts to disrupt the diel cycle by extension of light or dark periods or incident light reduction revealed that the growth periods observed under normal conditions were primarily associated with the diel cycle, and not a result of circadian rhythms. *A. anophagefferens* does encode a homolog of an animal-like cryptochrome containing a photolyase domain, meaning circadian responses to light are not necessarily absent in this system (Petersen et al., 2021). While circadian control of the cell cycle has only been studied in a few model species (*Chlamydomonas reinhardtii, Ostreococcus tauri, Phaeodactylum tricornutum*), it is possible that specific expression levels of cell cycle-associated genes are constrained to a circadian clock in *A. anophagefferens*, without strict constraint of carbon fixation and cell growth-associated genes (Coesel et al., 2009; Heijde et al., 2010; Beel et al., 2012; Petersen et al., 2021). This necessitates further analyses of transcriptomic and proteomic profiles of *A. anophagefferens* under a free-running clock to draw any further conclusions.

Pelagophytes like *A. anophagefferens* exist in open oceans at the deep chlorophyll maximum, spatially deeper than cyanobacteria and dinoflagellates (Latasa et al., 2017). Given a preference for decreased light and increased nutrient availability, it is possible that these cells are highly susceptible to photooxidative stress and DNA damage when unshaded (Latasa et al., 2017). If light stresses are a factor, cells may benefit from cell division and DNA synthesis occurring during dark periods. G1 to S phase likely requires a DNA damage checkpoint to be met, though the genes regulating this transition are not well defined in *A. anophagefferens* (Hlavová et al., 2011). It is possible that to proceed to the downstream transcriptional effects of the cell cycle, a photoreceptor-like trigger must first be deactivated, akin to the red/far-red phytochrome receptor in plants (Mawphlang and Kharshiing, 2017). This is supported by our work showing that *A. anophagefferens* cannot grow in 24-hour light.

### Virocells display arrested division phenotypes but continue to respond to light exposure

Infection of *A. anophagefferens* by *K. quantuckense* inhibited cell division during the dark period of the diel cycle. The virocells maintained the same size overnight, as opposed to the uninfected cells which decreased in average diameter in parallel with division. This may act as a benefit to virus propagation as increased volume of the cell may correspond to an ability to house more virus particles before lysis. Virocells also did not divide. It is possible that the cells were unable to enter mitosis either through degradation of the host genome by viral endonucleases or an increase in the density of early-stage viral particles. As well, if host microtubules were physically inhibited from correctly allocating to polar ends of the cell or chromosomal kinetochores by viruses packed into the host, it is feasible that the cell cycle could not progress. However, viral particle formation and the degradation of most organelles does not occur until later in the infection cycle. This leads to the possibility that transcription under viral infection prevents the cell cycle from progressing and locks the infected cells into a prolonged G2 phase before the cell can enter mitosis. This is notably not the only outcome of virus-host interaction, as infecting the partially resistant CCMP1850 with *K. quantuckense* reveals inhibition of division but consistent cycling of cell size (Gobler et al., 2007).

Another element of the infection dynamics of *A. anophagefferens* and *K. quantuckense* is the similar growth during the light cycle between infected and uninfected samples. As previous studies have shown, light is an important constraint on the burst size of *K. quantuckense* (Gann et al., 2020a). Combined with the fact that infection is not observed under zero light conditions, successful infection of a host may require a prolonged light. While transcription of viral genes begins within the first five minutes of infection, the cellular growth during light periods remains the same, meaning this portion of the growth cycle is perhaps relevant to the virus’s propagation. While we may define this period as nutritional stockpiling by an uninfected host to prepare for DNA synthesis and cell division, in the case of viral infection the same stockpiling must occur, only to be used in production of viral particles (Gann et al., 2020b). In considering what metabolic processes the virus alters for its own benefit, we also must consider which processes are left unaltered and how these too may serve a purpose in viral infection.

### *A. anophagefferens’* cell cycle is transcriptionally constrained to diel effects

Analysis of the viral infection transcriptome revealed that cell cycle arrest and regulation genes were changed during viral infection, but also that transcription of many cell cycle-associated genes in uninfected cells is constrained to a diel cycle. One of the most common observations of diel-driven differential expression was the expression of cohesin and condensin genes. The cohesin complex, which binds sister chromatids together following DNA replication and prior to anaphase, acts as an important intermediate complex before sister chromatids are segregated to opposite ends of the cell (Peters et al., 2008). The condensin complex begins functioning typically after the nuclear envelope has broken down from prophase to anaphase (Hirano, 2012; Leonard et al., 2015). Further research has shown that in *C. reinhardtii,* condensin subunits are likely involved in proper formation of the mitotic spindle (Breker et al., 2018). The expression patterns of these subunits, cohesin upregulated in the early night and condensin upregulated in the late night, suggest that cells were progressing through mitosis at these time points. Thus, DNA replication likely occurred either soon before or after the dark period began. Likewise, we have shown that cell division primarily occurs after 6-7 h in the dark (**Figure S3**), meaning condensin would be heavily expressed at this time point (Skibbens, 2019).

Further evidence for the temporal partitioning of cell cycle pathway genes is shown by the expression pattern of *pcna*. In the red alga *Cyanidioschyzon merolae*, *pcna* was used as a marker of cell cycle progression and peaked in fluorescence mid S-phase, before dissipating throughout the remainder of the cell cycle (Sumiya et al., 2014). A similar expression was observed in the transcriptomic dataset, with *pcna* exclusively peaking in expression in the early night. This indicates DNA synthesis may occur around the transition from light to dark. Along with *pcna*, several other genes including cyclin B (*ccnb2ii*), *cdc45* (which has been tied to replication fork initiation (Sanchez-Pulido and Ponting, 2011)), *cdc5*, five cohesin subunits, and the DNA damage repair gene *rrm2* show this pattern. We hypothesize that these genes are largely associated with DNA synthesis. Several genes expressed heavily in the early night are also detected at increased levels in the late day (*tp53i3*, *hrad1*, *atm,* and *erk*). Such genes can be attributed to acting as a catalyst for DNA damage repair and preventing cellular division with damaged or incompletely replicated DNA. Likewise, a close homolog to the yeast gene c*dc28*, which is active late in G1 phase (Mendenhall and Hodge, 1998), was upregulated in the early hours of the night, as well as mid-day, identifying an early trigger to encourage DNA synthesis. This gene is notably not expressed in the late night, signaling that many cells have progressed past the early stages of the cell cycle.

Among the genes expressed differentially in the late night were the p53 inhibiting *mdm2* homologs, *cdc20*, monopolar spindle 1 kinase (*mps1*), and two condensin subunits, of which *A. anophagefferens* only encodes three. Based on the presence of genes like *cdc20, mdm2* and *mps1* which actively either drive or regulate mitotic cell division (Mendoza et al., 2014; Pecani et al., 2022), it seems evident that the cells at the late time point were actively in the process of mitosis. Its expression in these contexts indicates cell division may be heavily down-regulated late in the day but encouraged after several hours in the night cycle.

### *K. quantuckense* infection drives transcriptional shifts in cell cycle regulation

Viral infection of *A. anophagefferens* revealed heavy regulation of genes associated with DNA replication. Several cohesin subunits and the *pcna* gene were upregulated late in the infection cycle after their transcript levels had decreased in the control samples. These genes may be involved in replication of the viral genome, which would justify such an expression level. Notably, many genes associated with cell cycle arrest based on DNA damage have altered expression levels during viral infection (**Figure 8**). Three homologs of *myt1/wee1* were downregulated under infecting conditions, one of which was downregulated at the three latter time points. WEE1 is typically considered a tumor suppressing protein and is often involved in the prevention of mitosis when DNA is damaged by way of phosphorylating CDK1 (Hlavová et al., 2011; Ghelli Luserna di Rorà et al., 2020). In accordance with this downshift in expression, four negative regulators of *wee1* (*cul1*, *skp1*, *rbx1*, and *prmt5*), were frequently upregulated during infection while three more DNA damage response genes (*rrm2*, *prkdc*, and *ddb2*) were downregulated. To further associate the DNA damage with viral genome replication, ubiquitination of PCNA by certain checkpoint proteins may lead to stalling of replication, thus an upregulation of this gene as well as a down-regulation of damage associated genes would allow for viral DNA synthesis to proceed (Moldovan et al., 2007).

**Figure 8.**
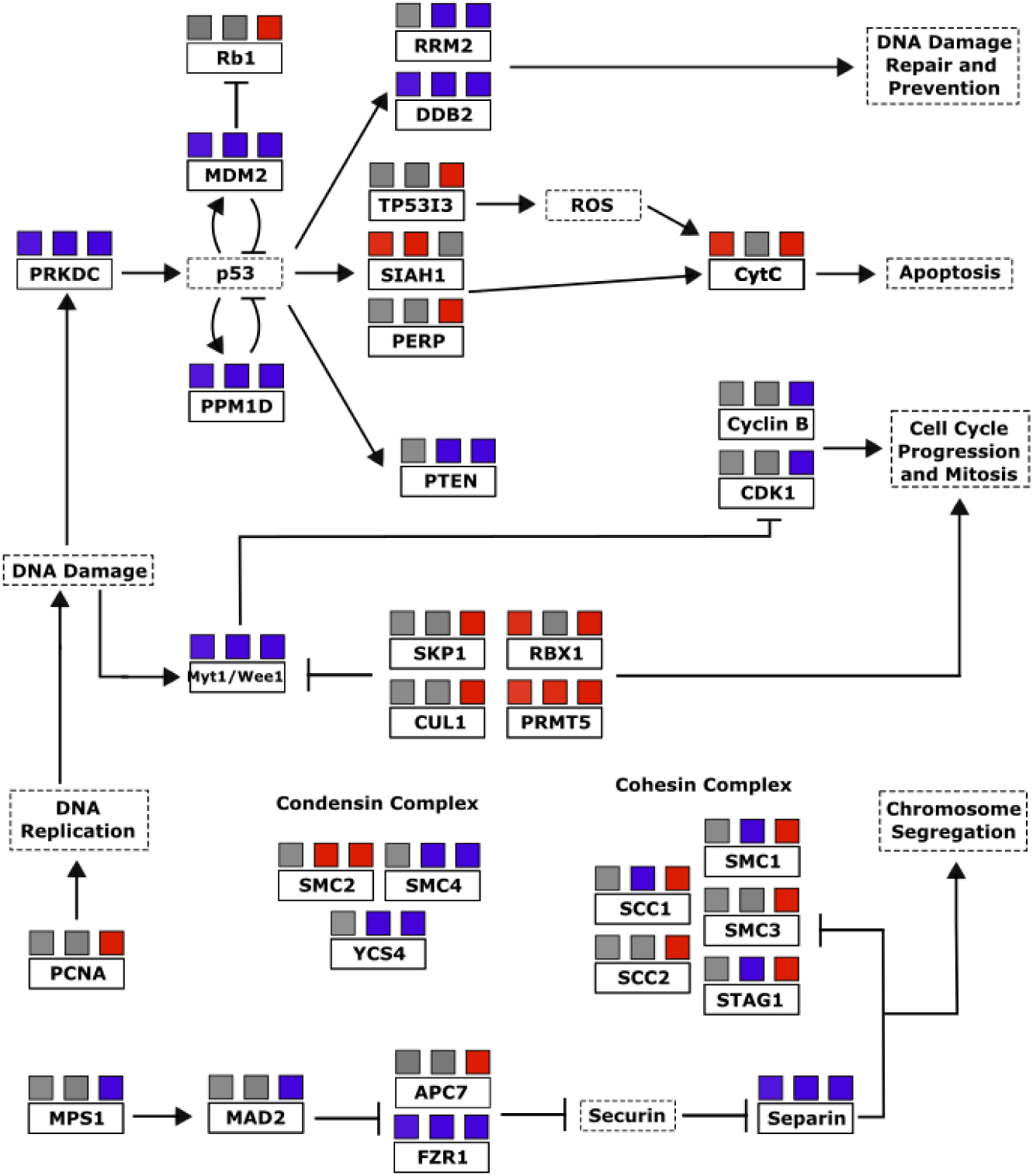
A simplified p53/cell cycle checkpoint pathway based on differential transcription of *A. anophagefferens* genes during viral infection. Genes present in the *A. anophagefferens* genome are indicated in boxes with solid outlines. Other genes not known to be encoded by *A. anophagefferens* or downstream effects of a certain pathway are in boxes with dashed outlines. Upregulation in samples is indicated by red boxes and downregulation is indicated by blue boxes, with the three boxes above each respective gene corresponding to their expression versus control samples during the late day, early night, and late night. Promotion of a gene/pathway is indicated by an arrow while inhibition is indicated by blunt arrows. Pathway is based on KEGG pathways map04110 and map04115.

From this concept there arises a conflicting dichotomy in the viral transcriptome in which genes that both lead to and prevent cell cycle arrest are regulated. While these DNA damage checkpoint genes may be downregulated, p53 inhibitors, which would naturally promote the cell cycle, are downregulated as well. We see a very complex expression pattern in which some of the downstream effectors of p53 like the apoptotic pathway and *rb1* are promoted under viral expression, while others that relate back to DNA damage control which might halt the cell cycle during S-phase are blocked in expression. Another perspective is that certain promoters of cell cycle progression are not altered by viral infection. While a total of 175 *A. anophagefferens* genes were functionally categorized as cell cycle genes, only 67 of them appeared in our differentially expressed dataset, meaning over 60% were statistically unchanged by viral infection. For example, *A. anophagefferens* encodes for 10 anaphase promoting complex genes, only one subunit was downregulated at one time point. While this does not confirm that these processes were not at all affected by the invading virus, it does suggest that continued activation of some of these pathways has at the very least a net neutral effect on viral particle production. A highly specialized virus must be able to streamline the infection process, and the host processes that are neutral, beneficial, or essential must remain active while anything deleterious be targeted for downregulation. For this reason, we believe that if the virus prevents the host cell from entering mitosis, the natural progression of the cell cycle may be important for the culmination of the infection cycle.

Though these possibilities are intriguing, our interpretations are limited by the lack of a complete cell cycle model in *A. anophagefferens*. Not only are most of the genes described herein attributed to putative functions through sequence homology, but many common genes also appear to be completely missing. For example, there are zero homologs to conventional CDK inhibitors encoded in any sequenced *A. anophagefferens* genome. Likewise, though we reference the p53-associated pathway for growth suppression often, there is no known homolog for p53 encoded by *A. anophagefferens* or any other plant or algal lineage. However, with such a robust representation of associated genes, including the direct downstream gene of *tp53i3* and the high abundance of *mdm2*-like p53 regulators, the presence of a functional p53 equivalent in *A. anophagefferens*, as well as other algae, is likely (Nedelcu, 2006).

### Export from the water column is enhanced by viral infection

Settling rate assessment of uninfected and infected *A. anophagefferens* helped link ecological relevance to physiological changes. While increases in cell size did not affect sinking velocity, viral infection increased vertical transport and suggests an increased rate of export from the water column. An interesting question that arises from the export of virally infected cells is whether the behavior is a result of viral activity inside the virocell or is instead host driven. One possibility was the increased production of viral capsid proteins increases density within the virocell, expediting sinking. Yet there was no significant increase in sinking velocity during later stages of infection (when virus proteins are more abundant within the cell) relative to the early (2 h) stage. A contrasting hypothesis is that infected *A. anophagefferens* cells are exported from the water column through metabolic shifts following infection. This would create a separation between uninfected cells and new viruses released into the water column (an innate defense against community infection). Moreover, given the inability of *K. quantuckense* to lyse *A. anophagefferens* in the dark, the expediated sinking could foster an opportunity for survival against infection, given cellular mechanisms a chance to purge the virus or enter a cyst-like state and prevent viral propagation (Ma et al., 2020). It is yet unclear if entering this resting stage has such an effect or if the lytic cycle continues once the cell is metabolically active again. This manifestation of the “virus shuttle” (Sullivan et al., 2017) provides a selective mechanism for reinforcement at evolutionary scales.

### Considerations for environmental sampling and diel cycles

This study illustrates the importance of sampling phototrophs throughout the solar day. Clear physiological and transcriptional differences were evident among *A. anophagefferens* cells depending on light history. Moreover viral transcripts in field samples have been tied to different stages of the diel cycle, with reads decreasing 10-fold between day samples and night samples on a viral species, but not genus level (Martinez-Hernandez et al., 2020) – to this end perhaps our observations should not be surprising. Yet this becomes an important caveat in the analysis of environmental-omics. Over 1,800 genes in uninfected cells were differentially expressed between at least two sampling points, corresponding to ∼11% of all predicted genes in *A. anophagefferens* (Gann et al., 2022). Such heterogeneity in gene expression among control samples is compelling and indicates that cells in non-synchronous infections are transcriptionally and metabolically distinct at different times of day. Yet some of the signal is also due to virus-shaped metabolism: if light shapes responses in the field as it has in our lab study, it means that when a significant portion of the population is infected (*e.g.,* up to 37.5%, Gastrich et al. 2004) that large degrees of variability in the data could simply be the infected *vs* non-infected state.

### Conclusions

We have demonstrated the effects the diel cycle on growth and division of the pelagophyte *A. anophagefferens* and that infection by a “giant virus”, *Kratosvirus quantuckense,* inhibits the diel cycling of cell size and cell division. Our findings demonstrate an important linkage between cellular energetics and physiology and that this process is interrupted by viral takeover. These findings also illustrate the importance of light in the infection cycle of viruses of phototrophic hosts. When considering the activity of marine viruses, it may become important to consider sampling multiple times throughout both the day and night to achieve a higher resolution on environmental viral infection. Likewise, ignoring virocells leaves large holes in measurements of ecological physiology. Further analysis into the transcriptome of individual cells (*i.e.*, single cell transcriptomics) in the presence and absence of infectious particles may better show the extent to which these cells vary throughout something as simple as the diel cycle. Still, our findings have revealed a continuously altering physiological profile in algae which likely extends beyond light into other environmental stimuli that should be further explored on an *in situ* basis.

## Supporting information

Supplemental Tables, Images and Methods

## Conflict of Interest

The authors declare that the research was conducted in the absence of any commercial or financial relationships that could be construed as a potential conflict of interest.

## Author Contributions

**ART:** Conceptualization, Formal analysis, Investigation, Writing – original draft, writing – review and editing; **EEC:** Conceptualization, Formal analysis, Investigation, Writing – original draft, writing – review and editing; **ARS**: Investigation, writing – review and editing; **SWW:** Conceptualization, Supervision, writing – review and editing.

**Funding:** This work was supported by the Simons Foundation (735077) and funds from the National Science Foundation (IOS1922958). ARS was supported by a REU Site Award (NSF 2050743).

## Acknowledgements

We thank Dr. Brittany Zepernick, Dr. Naomi Gilbert, Dr. Robbie Martin, Dr Gary LeCleir, Laura Smith, and David Niknejad for their contributions and valuable discussions.

## Notes

### Competing Interest Statement

The authors have declared no competing interest.

