## Supplemental Tables, Images and Methods for "The diel disconnect between cell growth and division in *Aureococcus* is interrupted by giant virus infection"

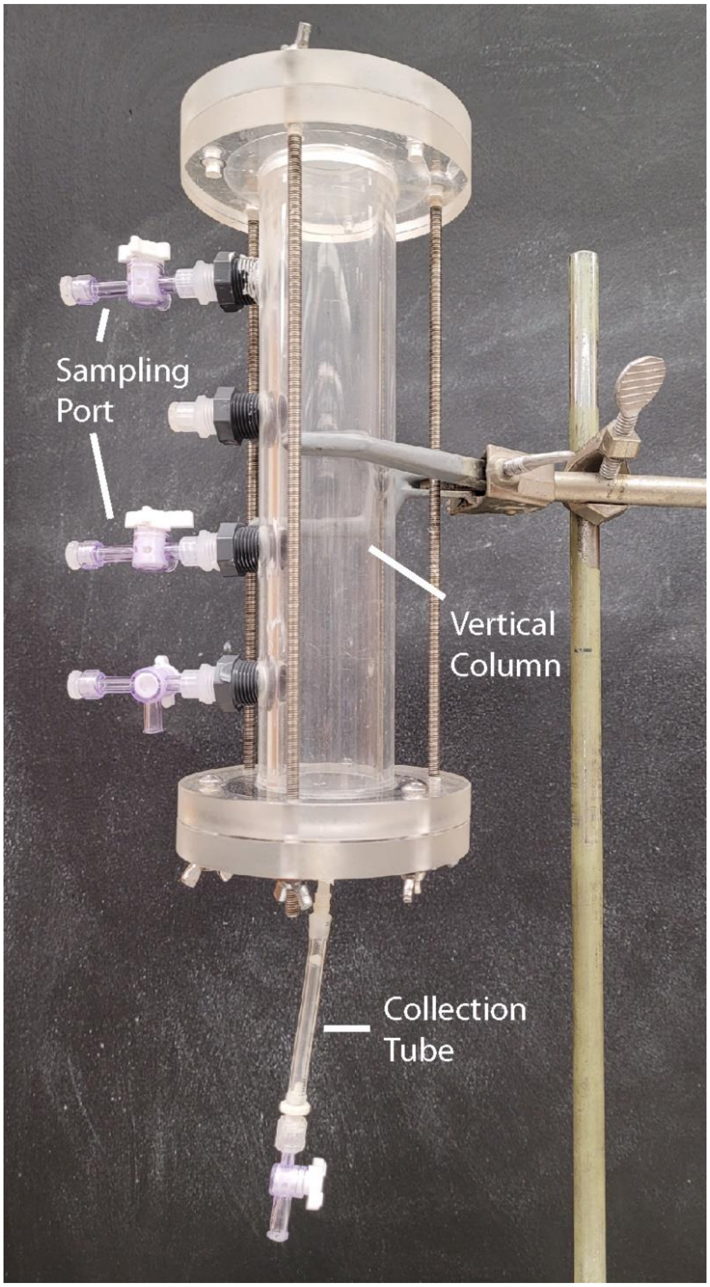


**Figure S1.** The setup of the vertical sinking column used in this study.

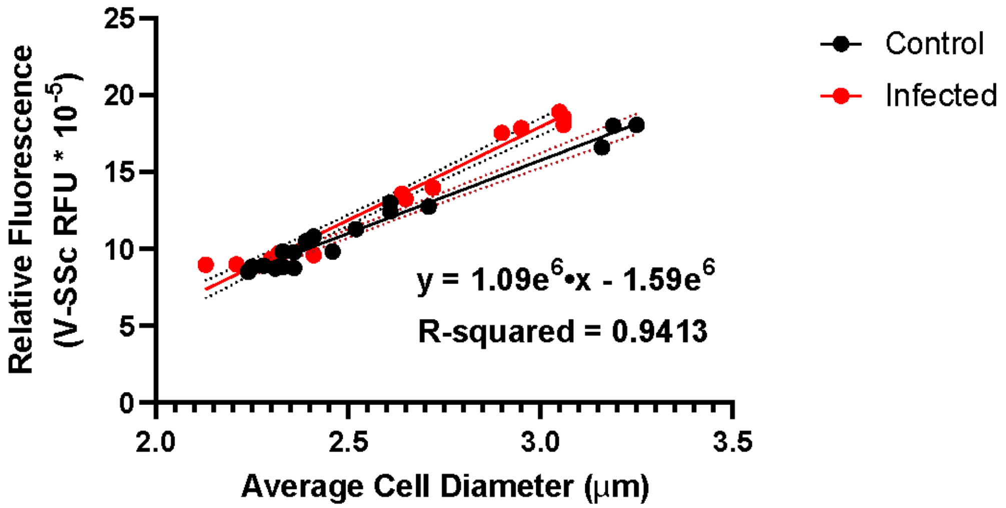


**Figure S2.** Correlation between violet side-scatter measurements taken on the Cytoflex flow cytometer and average cell diameter measured *via* FlowCam.

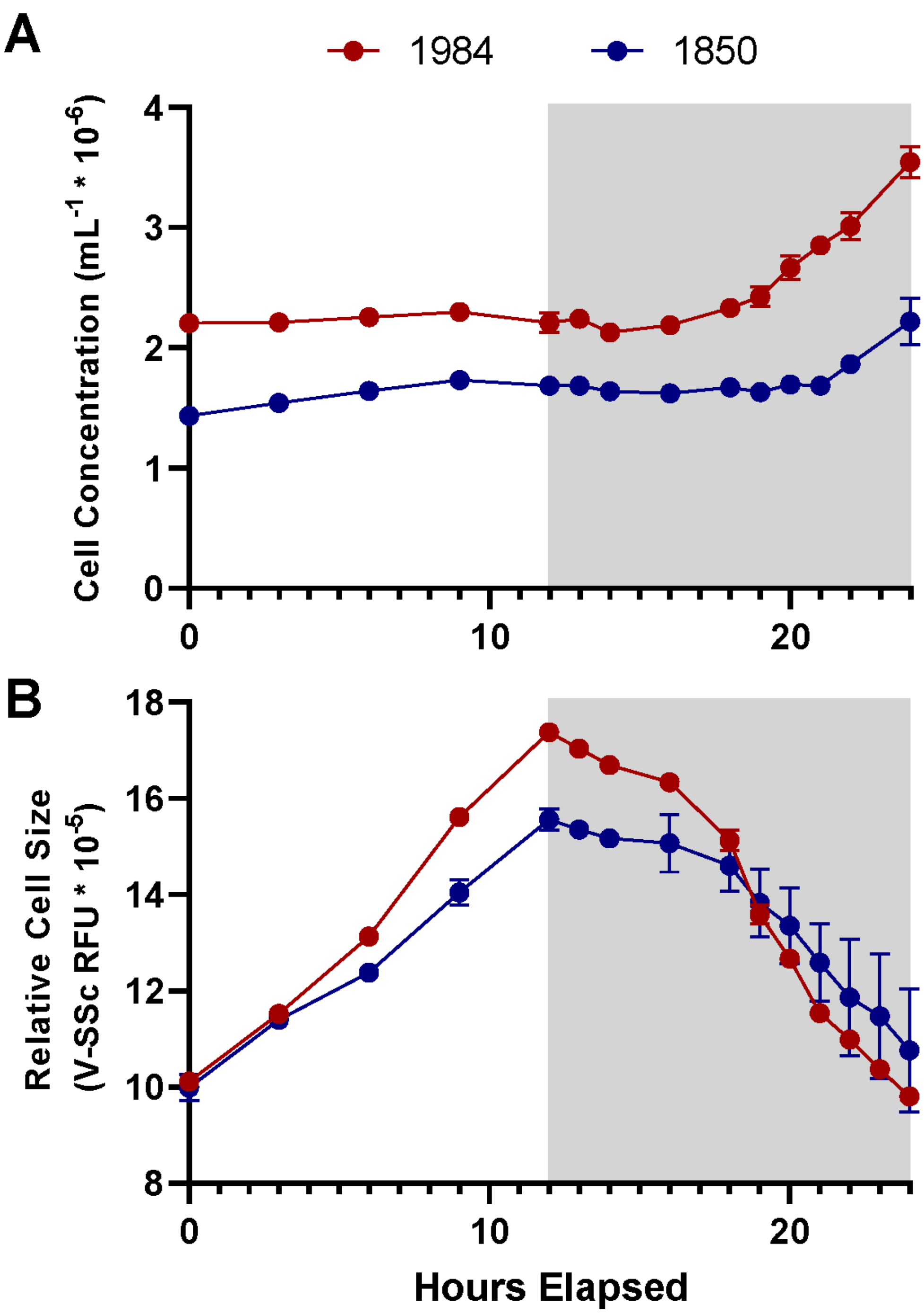


**Figure S3.** Cell density (A) and approximate cell diameter (B) of two *A. anophagefferens* strains, CCMP1984 (red) and CCMP1850 (blue) with particular attention paid to the night period.

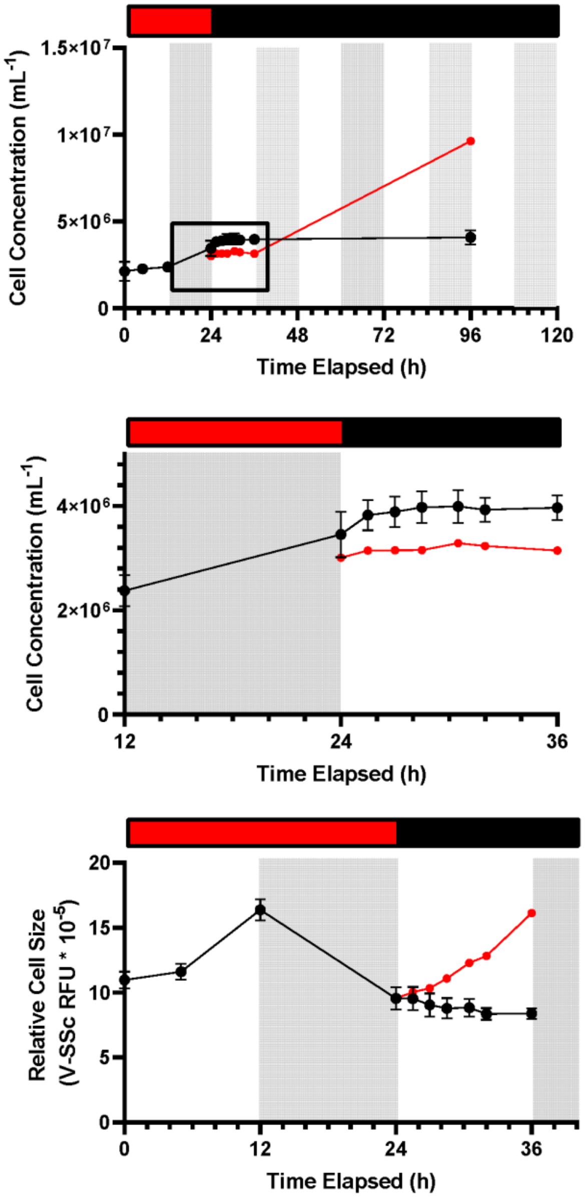


**Figure S4.** Free-running clock experiments on *A. anophagefferens*. Cultures were entrained to a consistent diel cycle before and either left in the same cycle at the start of the light period (red) or moved to a parallel fully dark incubator (black) at the 24-hour timepoint. Cell concentrations were measured frequently in the first 12 hours of the free-running clock (A) after 96 hours (B) either in complete darkness or in the regular diel cycle. Cell diameter was also measured within the first 12 hours of the free-running clock (C). Grey and white backgrounds only refer to the light cycle of the entrained samples while red and black periods above the graphs refer to the entrainment period and the free-running clock.

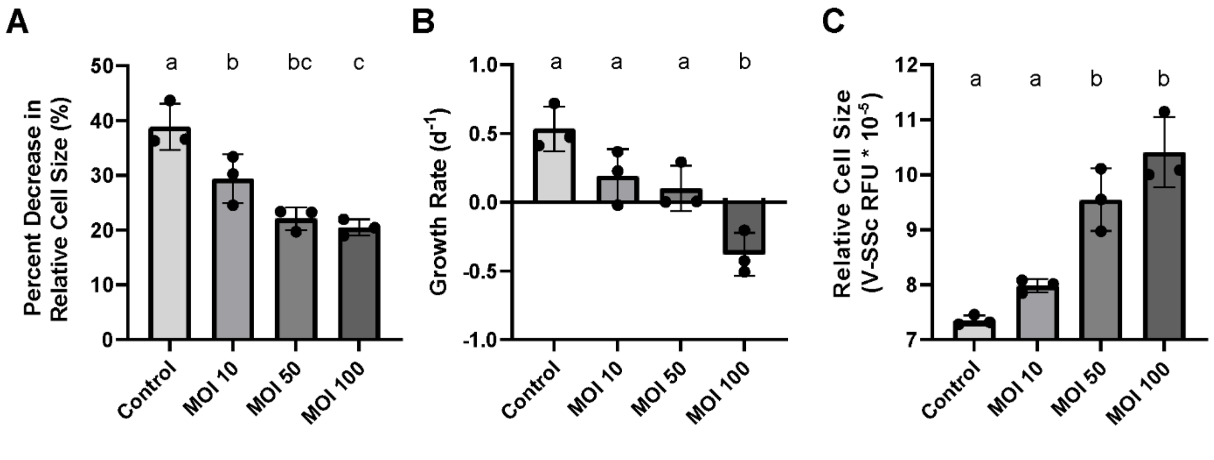


**Figure S5.** (A) Effect of decreasing M.O.I. on the average cell diameter in cultures 24 hours after infection. (B-E).

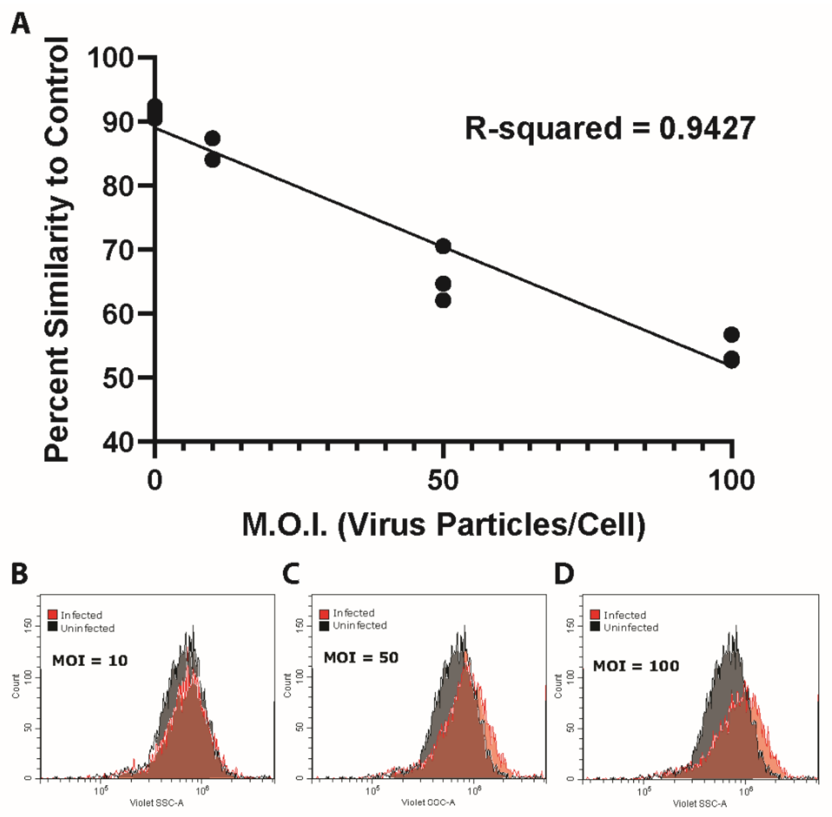


**Figure S6.** Percent similarity between control samples and infected treatment samples were calculated *via* the overlay function in the Cytoflex software CytExpert (A). Correlation between percent similarity to the control after 23 hours of infection and MOI is denoted (n = 3). Overlap of individual control samples and samples of each MOI are shown (B-D).

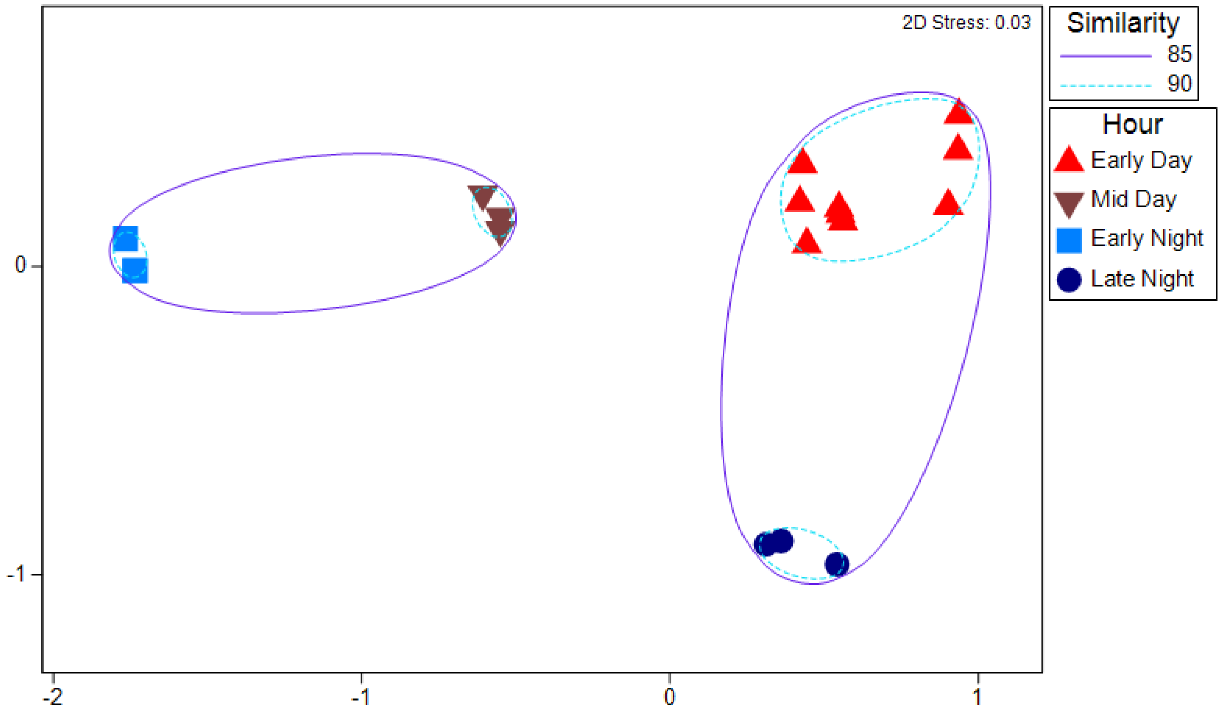


**Figure S7.** NMDS analysis of all uninfected samples colored based on the time of day they were taken.

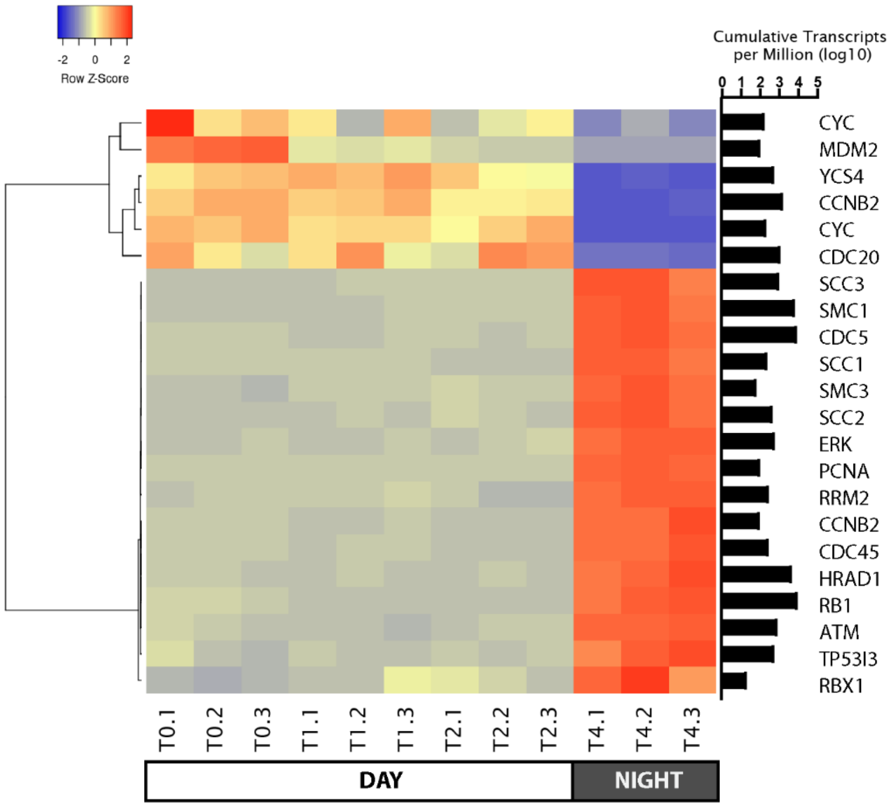


**Figure S8.** Heatmap of DESeq2 normalized cell cycle gene read counts that were found to be differentially expressed (p-value < 0.05, log2fold change > 2 or < -2) between the early day and the early night control transcriptome samples. The sum of transcripts across all treatments for each individual gene are indicated in log(TPM).

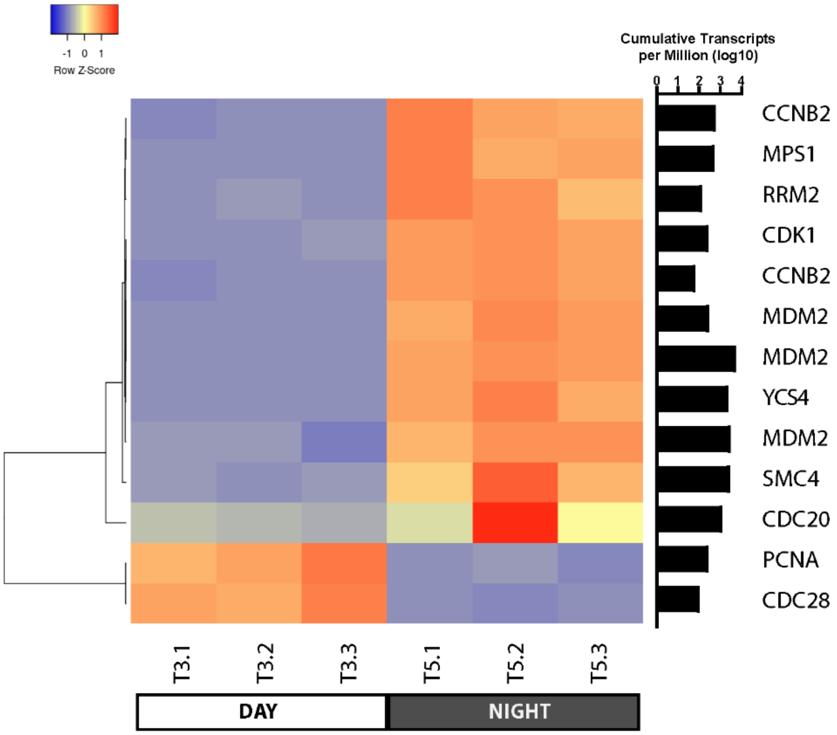


**Figure S9.** Heatmap of DESeq2 normalized cell cycle gene read counts that were found to be differentially expressed (p-value < 0.05, log2fold change > 2 or < -2) between the late day and late night control transcriptome samples. The sum of transcripts across all treatments for each individual gene are indicated in log(TPM).

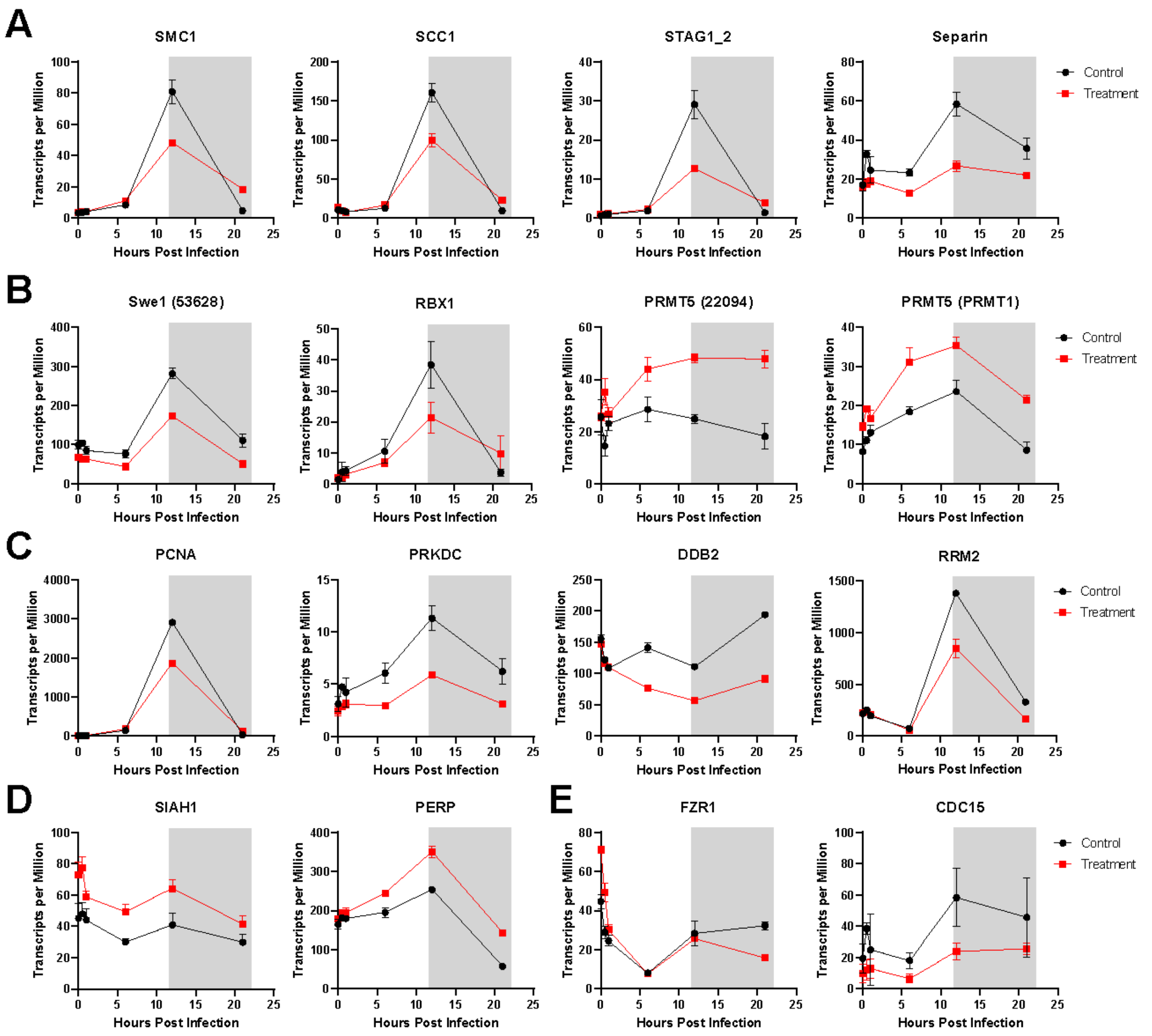


**Figure S10.** A selection of differentially expressed genes between and their relative expression patterns (TPM) throughout the infection cycle between control uninfected (black) and infected (red) treatments. (A) Expression of cohesin associated genes, (B) Swe1 and several of its regulatory genes, (C) genes involved in DNA replication and regulation of DNA damage, (D) apoptosis associated genes, and (E) drivers of mitotic division.

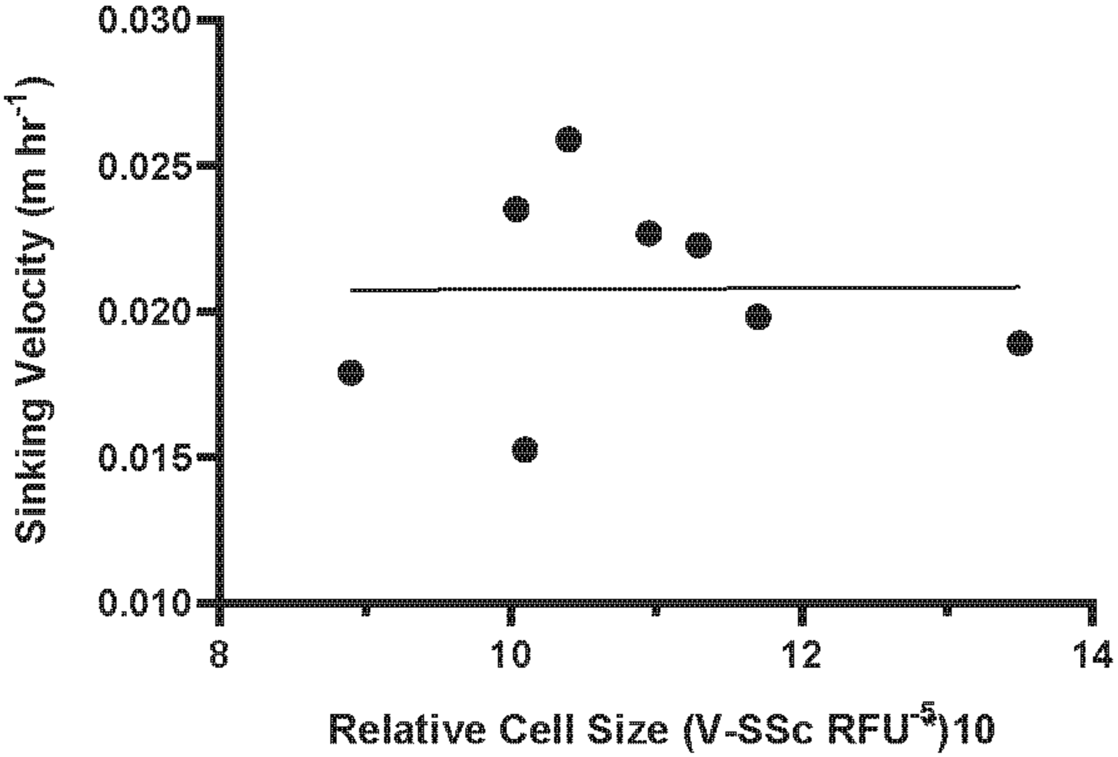


**Figure S11**. Correlation between relative fluorescence (V-SSc) and sinking velocity of *A. anophagefferens* CCMP1984 cells.

**Table S1.** Growth dynamics of *A. anophagefferens* in the presence of either total light or total dark. Rates are given ± S.D.

| | CCMP1707 | | CCMP1850 | | CCMP1984 | |
|---|---|---|---|---|---|---|
| | **Dark** | **Light** | **Dark** | **Light** | **Dark** | **Light** |
| **Growth Rate @24 hr ($d^{-1}$)** | 0.016 (0.001) | -0.095 (0.077) | 0.029 (0.028) | -0.118 (0.017) | 0.002 (0.020) | -0.021 (0.028) |
| **Growth Rate @48 hr ($d^{-1}$)** | 0.012 (0.020) | -0.147 (0.116) | 0.022 (0.021) | -0.094 (0.016) | 0.007 (0.013) | 0.017 (0.055) |
| **% Change in Cell Size (@24 hr)** | -0.083 (0.024) | 0.275 (0.048) | -0.121 (0.004) | 0.144 (0.020) | -0.144 (0.025) | 0.381 (0.054) |
| **% Change in Cell Size (@48 hr)** | -0.211 (0.033) | 0.407 (0.099) | -0.201 (0.027) | 0.11 (0.012) | -0.238 (0.016) | 0.537 (0.089) |

**Table S2.** All *A. anophagefferens* cell-cycle associated genes which are either upregulated or downregulated 1.5-fold (p-value < 0.05) at either the 6-hour, 12-hour, or 24-hour timepoints in the presence of *K. quantuckense.*

| Gene_ID | Gene | 6-hour (log2-fold change) | 12-hour (log2-fold change) | 21-hour (log2-fold change) | Notes |
|---|---|---|---|---|---|
| **Cohesin, Condensin, and Other Associated Genes** | | | | | |
| AURANDRAFT_64106 | SCC1 | NA | Down (-0.64) | Up (1.14) | *Cohesin complex subunit 1* |
| AURANDRAFT_70503 | SMC1 | NA | Down (-0.69) | Up (1.81) | *Structural maintenance of chromosome 1; Cohesin subunit* |
| AURANDRAFT_72635 | SMC3 | NA | NA | Up (1.55) | *Structural maintenance of chromosome 3; Cohesin subunit* |
| AURANDRAFT_60440 | STAG1_2 | NA | Down (-1.15) | Up (1.34) | *Cohesin complex subunit SA-1* |
| AURANDRAFT_26480 | SCC2 | Down (-0.94) | Down (-1.49) | Down (-1.39) | *Cohesin loading factor subunit 2* |
| AURANDRAFT_63990 | SCC2 | NA | NA | Up (1.43) | *Cohesin loading factor subunit 2* |
| AURANDRAFT_36910 | SMC2 | NA | Up (0.75) | Up (1.84) | *Structural maintenance of chromosome 2; Condensin subunit* |
| AURANDRAFT_72033 | SMC4 | NA | Down (-1.02) | Down (-1.50) | *Structural maintenance of chromosome 4; Condensin subunit* |
| AURANDRAFT_65310 | YCS4 | NA | Down (-0.80) | Down (-1.44) | *Condensin complex subunit 1* |
| AURANDRAFT_72664 | ESP1;NDC80 | Down (-0.89) | Down (1.07) | Down (-0.87) | *Separin* |
| **Swe1 Regulation** | | | | | |
| AURANDRAFT_53628 | Myt1;Wee1 | Down (-0.84) | Down (-0.65) | Down (-1.30) | *Mitosis inhibitor protein kinase* |
| AURANDRAFT_71641 | Myt1;Wee1;Swe1 | NA | Down (-0.92) | Down (-1.31) | *Mitosis inhibitor protein kinase* |
| AURANDRAFT_72516 | Myt1;Wee1;Swe1 | NA | Down (-0.94) | Down (-1.75) | *Mitosis inhibitor protein kinase* |
| AURANDRAFT_22094 | PRMT5 | Up (0.60) | Up (1.01) | Up (1.24) | *Type II protein arginine methyltransferase; Swe1 regulation* |
| PRMT1 | PRMT5 | Up (0.74) | Up (0.64) | Up (1.16) | *Type II protein arginine methyltransferase; Swe1 regulation* |
| AURANDRAFT_22764 | RBX1 | Up (0.79) | NA | Up (1.19) | *E3 ubiquitin-protein ligase; Swe1 regulation* |
| AURANDRAFT_8889 | RBX1 | NA | Down (-0.80) | Up (1.31) | *E3 ubiquitin-protein ligase; Swe1 regulation* |
| AURANDRAFT_58667 | SKP1 | NA | NA | Up (1.82) | *S-phase kinase-associated protein 1; Swe1 regulation* |
| AURANDRAFT_52007 | CUL1 | NA | NA | Up (0.80) | *Cullin 1; Swe1 regulation* |
| AURANDRAFT_55274 | CUL1 | NA | Down (-0.58) | NA | *Cullin 1; Swe1 regulation* |
| AURANDRAFT_4905 | CDC5 | NA | Down (-1.17) | NA | *Cell division control protein 5; Swe1 regulation* |

**Table S2 Continued**

**Apoptotis Associated Genes**

| | | | | | |
|---|---|---|---|---|---|
| AURANDRAFT_28360 | CYC | NA | Down (-0.67) | NA | *Cytochrome C* |
| AURANDRAFT_59926 | CYC | Up (0.72) | NA | Up (1.41) | *Cytochrome C* |
| AURANDRAFT_31918 | PERP | NA | NA | Up (1.16) | *TP53 apoptosis effector proteins* |
| AURANDRAFT_28417 | TP53I3 | NA | NA | Up (0.65) | *P53-inducible protein 3* |
| AURANDRAFT_20166 | SIAH1 | Up (0.69) | Up (0.71) | NA | *E3 ubiquitin-protein ligase; methyltransferase domain* |
| AURANDRAFT_72557 | LRDD | NA | NA | Down (-0.97) | *Death domain-containing protein* |
| **p53 Negative Feedback** | | | | | |
| AURANDRAFT_12504 | MDM2 | Down (-1.23) | Up (2.79) | NA | *E3 ubiquitin-protein ligase; p53 regulation* |
| AURANDRAFT_24297 | MDM2 | Down (-0.97) | Down (-1.72) | Down (-1.03) | *E3 ubiquitin-protein ligase; p53 regulation* |
| AURANDRAFT_27943 | MDM2 | NA | Down (-1.18) | Down (-0.74) | *E3 ubiquitin-protein ligase; p53 regulation* |
| AURANDRAFT_3154 | MDM2 | Down (-0.85) | Down (-1.51) | Down (-1.03) | *E3 ubiquitin-protein ligase; p53 regulation* |
| AURANDRAFT_33333 | MDM2 | NA | Down (-0.89) | NA | *E3 ubiquitin-protein ligase; p53 regulation* |
| AURANDRAFT_72197 | MDM2 | NA | Down (-0.63) | NA | *E3 ubiquitin-protein ligase; p53 regulation* |
| AURANDRAFT_63280 | PPM1D | Down (-0.77) | Down (-1.00) | Down (-1.08) | *Protein phosphatase 1D; p53 regulation* |
| **Regulation of DNA Replication** | | | | | |
| AURANDRAFT_70336 | MCM3 | NA | Down (-0.79) | NA | *DNA replication licensing factor 3* |
| AURANDRAFT_63773 | MCM4 | Up (0.71) | NA | NA | *DNA replication licensing factor 4* |
| AURANDRAFT_33815 | MCM4 | NA | Down (-0.60) | NA | *DNA replication licensing factor 4* |
| AURANDRAFT_38450 | MCM5 | NA | Down (-0.75) | Down (-0.68) | *DNA replication licensing factor 5* |
| AURANDRAFT_24050 | MCM6 | NA | Down (-0.59) | NA | *DNA replication licensing factor 6* |
| AURANDRAFT_53352 | MCM7 | NA | Down (-0.84) | Down (-0.81) | *DNA replication licensing factor 7* |
| AURANDRAFT_72178 | MCM7 | NA | Down (-0.85) | NA | *DNA replication licensing factor 7* |
| AURANDRAFT_63471 | ORC2 | NA | Down (-0.67) | Down (-0.62) | *Origin recognition complex subunit 1* |
| AURANDRAFT_70163 | PCNA | NA | NA | Up (1.77) | *Proliferating cell nuclear antigen* |
| AURANDRAFT_62055 | CDC6 | NA | NA | Down (-1.17) | *Cell division control protein; DNA replication-associated* |
| AURANDRAFT_20277 | CDC7 | NA | NA | Up (0.62) | *Cell division control protein; DNA replication-associated* |
| AURANDRAFT_71439 | DDB2 | Down (-0.91) | Down (-0.91) | Down (-1.25) | *DNA damage-binding protein 2* |
| AURANDRAFT_68528 | PRKDC | Down (-1.09) | Down (-0.90) | Down (-1.18) | *DNA-dependent protein kinase* |

| | | | | | |
|---|---|---|---|---|---|
| RRS3 | RRM2 | NA | Down (-0.64) | Down (-1.18) | *Ribonucleoside-diphosphate reductase M2* |
| AURANDRAFT_64915 | CCDN2 | NA | Up (0.58) | NA | *G1/S-specific cyclin D2* |
| **Regulation/Termination of Mitosis** | | | | | |
| AURANDRAFT_22946 | APC7 | NA | NA | Up (0.61) | *Anaphase-promoting complex subunit 7* |
| AURANDRAFT_28025 | APC7 | NA | NA | Up (0.62) | *Anaphase-promoting complex subunit 7* |
| AURANDRAFT_28134 | APC7 | NA | Down (-0.63) | NA | *Anaphase-promoting complex subunit 7* |
| AURANDRAFT_9374 | APC11 | Up (0.77) | NA | NA | *Anaphase-promoting complex subunit 11* |
| AURANDRAFT_28425 | FZR1 | NA | NA | Down (-1.18) | *Cell division cycle 20 protein; cofactor of APC complex* |
| AURANDRAFT_12473 | CCNB2 | NA | NA | Down (-0.94) | *G2/mitotic-specific cyclin B2* |
| AURANDRAFT_38949 | CCNB2 | NA | NA | Down (-0.75) | *G2/mitotic-specific cyclin B2* |
| AURANDRAFT_64231 | CCNB2 | NA | NA | Down (-1.37) | *G2/mitotic-specific cyclin B2* |
| AURANDRAFT_71534 | CCNB2 | NA | NA | Down (-0.59) | *G2/mitotic-specific cyclin B2* |
| AURANDRAFT_31094 | CDK1 | NA | NA | Down (-0.97) | *Cyclin-dependent kinase 1; G2 to M-phase* |
| AURANDRAFT_13918 | TTK/MPS1 | NA | NA | Down (-0.88) | *Monopolar spindle kinase* |
| AURANDRAFT_19390 | MAD2 | NA | NA | Down (-0.75) | *Mitotic spindle assembly checkpoint protein* |
| AURANDRAFT_70730 | HRAD1 | NA | NA | Up (0.88) | *Cell cycle checkpoint protein* |
| AURANDRAFT_71576 | PTEN | NA | Down (-0.95) | Down (-1.23) | *Phosphatase and tensin homolog; tumor supression* |
| AURANDRAFT_32347 | TEM1 | NA | NA | Up (1.07) | *GTP-binding protein; termination of M-phase* |
| AURANDRAFT_32157 | CDC15 | Down (-1.51) | Down (-1.23) | NA | *Protein kinase; cell division control protein* |
| **Other Cell Cycle-Associated Genes** | | | | | |
| AURANDRAFT_30303 | ERK;MAPK1_3 | NA | NA | Down (-1.09) | *Mitogen-activated protein kinase* |
| AURANDRAFT_31374 | ERK/MAPK1_3 | NA | NA | Up (0.79) | *Mitogen-activated protein kinase* |
| AURANDRAFT_39527 | ERK/MAPK1_3 | NA | NA | Down (-0.63) | *Mitogen-activated protein kinase* |
| AURANDRAFT_55239 | ERK/MAPK1_3 | Up (0.59) | NA | Up (1.21) | *Mitogen-activated protein kinase* |
| CDK1 | CDC28/CDC2 | NA | Down (-0.73) | NA | *Cyclin-dependent kinase* |
| AURANDRAFT_58854 | CDK7 | NA | NA | Up (0.87) | *Cyclin-dependent kinase 7* |
| AURANDRAFT_69594 | CCNH | NA | NA | Up (0.67) | *Cyclin H* |
| AURANDRAFT_12637 | RB1 | NA | NA | Up (1.24) | *Retinoblastoma-associated protein-like; ribonuclease H2* |
| AURANDRAFT_68494 | GSK3B | Down (-0.97) | NA | NA | *Glycogen synthase kinase 3* |
